# Too much or not enough? Optimal level of human intracranial theta activity for rule-switching in the subthalamo-medio-prefrontal circuit

**DOI:** 10.1101/2023.12.11.571129

**Authors:** Maëva Laquitaine, Mircea Polosan, Philippe Kahane, Stephan Chabardes, Jérôme Yelnik, Sara Fernandez-Vidal, Philippe Domenech, Julien Bastin

**Author notes:** These authors contributed equally (Philippe Domenech, Julien Bastin).

## Abstract

The ability to strategically switch between rules associating stimuli and responses as a function of changing environmental demands critically depend on a neural circuit including the dorsomedial prefrontal cortex (dmPFC) and the basal ganglia. However, the precise neural implementations of rule switching remain unclear. To address this issue, we recorded local field potentials from two groups of rare patients performing a rule-switching paradigm: (1) deep brain recordings of the subthalamic nucleus (STN) in patients with obsessive-compulsive disorder, and (2) stereo-electroencephalogram from dmPFC of drug-resistant epileptic patients. We fitted a hierarchical drift-diffusion model (HDDM) to patients’ choice behavior and found that rule-switching was associated with a shift in the starting point of evidence accumulation (*z*), effectively disentangling rule switches from the selection of a new response. At the neural level, we found that theta band (5-10 Hz) activity increased in dmPFC and STN during switch compared to non-switch trials, while temporally delayed and excessive levels of theta activity led to premature switch errors. This seemingly opposing impact of increased theta rhythms in successful and unsuccessful switching could be explained mechanistically using a neural HDDM, as trial-by-trial fluctuations in theta power negatively correlated with the subjects’ starting point parameter. Together, these results shed a new light on the neural mechanisms underlying the rapid reconfiguration of stimulus-response associations, revealing a Goldilocks’ effect of theta band activity on rule switching behavior.

## Introduction

Rule switching allows us to rapidly adapt to changing demands and depends on a set of high-level cognitive functions from the anticipated application of abstract rules and the inhibition of no longer relevant choices to the selection and execution of the most adaptive response^1,2^. However, the neuro-computational mechanisms underlying our ability to rapidly adjust our behavior in response to unpredictable and sudden environmental changes remains poorly understood.

Early evidence suggests that the hyperdirect pathway connecting the dorso-medial prefrontal cortex (dmPFC) to the subthalamic nucleus (STN) could be related to rule-switching, action stopping or response conflict^3–6^. Yet, previous studies have mainly focused on action stopping or speed-accuracy trade-off adjustments to “conflict” (also referred to as choice difficulty or choice uncertainty)^7–10^. Previous human intracranial studies showed that human dmPFC and STN neurons’ firing rate, as well as theta (∼5-10 Hz) prefronto-subthalamic activity, increase when the best response is more difficult to select and require longer decision time^10–14^. Conversely, action stopping increases both neurons firing rate and beta-band activity (∼15-30 Hz) in the same dmPFC-STN circuit^15–22^. By contrast, non-invasive electrophysiological studies of task switching yield mixed findings in terms of temporal dynamics, frequency regime and putative brain circuits^23–25^, highlighting the need for more direct human electrophysiological recordings to establish a precise mapping between prefronto-subthalamic neural dynamics and key cognitive processes underlying task-switching.

Drift diffusion models posit that response selection proceeds by accumulating evidence up to a decision threshold in a winner take all fashion. This class of model was also shown to be relevant in the context of rule switching to tease apart the behavioral variability arising from switching between set of rules and the selection of a new response, which seems to occur sequentially^26,27^. Conversely, studies combining direct electrophysiological recordings with a drift diffusion modeling of perceptual choices revealed that decision-threshold adjustments to difficulty were associated with STN-dmPFC neural activity either in theta or beta bands, depending on the cognitive context^9,28^. However, to the best of our knowledge, such a neuro-computational dissection has never been performed using invasive human electrophysiological recordings in the dmPFC-STN circuitry during rule switching. Moreover, the use of different experimental tasks in human neuroimaging and monkey single-cell literatures has made it difficult to compare the cognitive and neural processes at play during rule switch across techniques and species (but see^38^ for an exception).

Thus, critical questions about how neural activity relates to rule switching remain unaddressed: Do nodes in the dmPFC-STN hyperdirect circuit exhibit specific temporo-frequency signatures associated with switching between sets of rules? Does neural activity within these brain regions relate to trial-by-trial fluctuations in behavioral performance, suggesting a close link between successful neural computations and participants’ ability to implement rule switching? What are the neuro-computational principles underlying these brain-behavior associations?

To address these questions, we used intracerebral recordings collected in two groups of rare neuropsychiatric patients implanted with depth electrodes, which allowed us to record from the dmPFC and the STN (respectively), while these patients performed a rule-switching paradigm previously used in monkeys^29,30^ to record multiunit activity from both the dmPFC and the STN. We show that behavioral switching depends on theta activity occurring in both dmPFC and STN. Critically, trial-by-trial fluctuations of theta activity negatively correlated with task switching performance in both brain structures, which translated in a neural HDDM accounting for observed RTs and performances as a modulation of the initial level of evidence by theta activity, prior to response selection, during switch trials. Our findings demonstrate that although theta activity in the dmPFC and in the STN increases on average during switch trials, too high an increase might paradoxically induce premature incorrect switch responses that are detrimental to adaptation.

## Results

We administered a reactive rule-switching paradigm to two groups of post-surgery neuropsychiatric patients while intracranial electroencephalographic activity (iEEG) was recorded and to one group of healthy controls: (1) the first group consisted of three patients with drug-resistant partial epilepsy implanted with linear depth macroelectrodes in the dorso-medial prefrontal cortex (dmPFC), (2) the second group consisted of four patients with severe and drug-resistant OCD implanted with deep brain stimulation macroelectrodes in the subthalamic nucleus (STN), and (3) the third group consisted of ten healthy controls (from whom we collected only behavioral recordings). We report demographic and clinical data in Table S1-2.

Each trial started with two colored squares (yellow/pink) randomly displayed on each side of a central white square (**Fig. 1**). After 500ms, the white central square turned either pink or yellow (ongoing rule) to prompt the patient to press a button indicating on which side of the screen the cue matching the central square color was displayed (left/right bimanual response). This stimulus-response association rule pseudo-randomly changed every 2 to 6 trials. Overall, the task consisted of two to four sessions composed of 50 switch trials each (138 ± 49 switch trials per subject). Patients were instructed to select the correct response as quickly and as accurately as they could. Hence, the task required to form a preliminary response by preemptively applying the previous rule as soon as the current stimulus pair was displayed. In switch trials (20% of trials), patients had to override this preliminary response to instantiate and apply the new rule, resulting in the selection of the alternative action.

**Figure 1.**
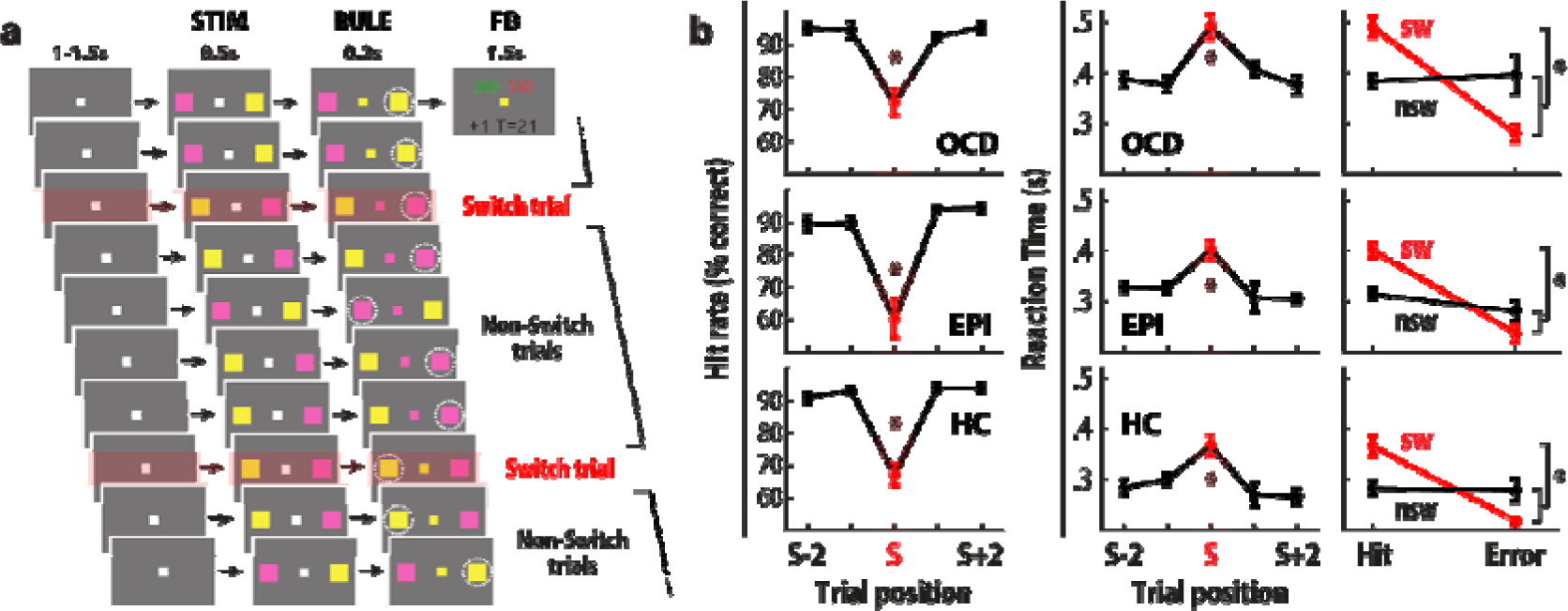
Behavioral task and results. (a) Task-switching paradigm. Patients had to indicate on which side the stimulus (STIM) matching the central square color (RULE) was. This response was followed by a feedback cue (FB) indicating whether it was correct and/or whether it fell within the allowed response time window. **(b) Behavioral performances of patients with OCD (upper row; n=4), epilepsy (EPI: middle row; n=3) and of healthy controls (HC: bottom row; n=10).** Evolution of hit rates and reaction times averaged according to trial relative position to switch trials (left and central panels) or as a function of trial type (red: switch; black: non-switch) and accuracy (hit or error). Error-bars indicate SEM between patients.

To promote speeded decisions, in line with the original work by Isoda et al. (2007), we implemented a response time window for non-switch trials (excluding switch trials, which typically have longer durations). An incorrect feedback was given during non-switch trials whenever the response time exceeded a predefined limit. The limit was adjusted based on participants’ performance after each switch trial: it increased by a 40 ms increment after incorrect switch trials and decreased by the same increment after correct switch trials, ensuring adaptive stringency. Note that this procedure did not censor observed RT distributions and maintained a consistent level of time pressure throughout the task as participants quickly reached a (fixed) floor value (300 ms).

### Behavioral performance: the cost of switching

Given the striking similarity between OCD and epileptic patients’ behavioral patterns (see Fig. 1), in the following, we report behavioral statistics pooled across the two groups of patients. It is noteworthy that these behavioral patterns closely mirrored those observed in healthy controls (Fig. 1b), while we also found consistent findings from ANOVAs followed by post-hoc tests performed separately for each group and even for each patient within the group (dmPFC group: Fig. S1; STN group: Fig. S2; see also Table S3 for statistics). First, we performed an analysis of variance (ANOVA) on mean reaction times (RTs), which demonstrated a significant interaction between trial type (switch vs. non-switch) and accuracy (hit vs. error) (F_(1,24)_ = 15.12; P<10^-3^; Fig. 1b; repeated measures ANOVA). Post-hoc comparisons confirmed that RTs on correct trials were significantly longer on switch trials in comparison to non-switch trials (99 ± 11 ms; t_(6)_= 9.14; P <10^-3^; paired two-tailed Student’s t test). RTs were also significantly faster during incorrect switch trials relative to incorrect non-switch trials, supporting the idea that errors during switch trials resulted from a failure to override the preliminary response to apply the new rule (-86 ms ± 28 ms; t_(6)_= -3.07; P <0.05; Fig. 1b; paired two-tailed Student’s t test). Unsurprisingly, patients also made significantly more errors on switch trials in comparison to non-switch trials (switch cost_(error)_= 26 ± 3 %; t_(6)_= 8.09; P <10^-3;^ paired two-tailed Student’s t test).

### Rule-switching: Neural activity in the dmPFC-STN network

To investigate whether neural activity in the dmPFC-STN network reflected rule-switching, we first contrasted the power estimated in the time-frequency domain between correct switch (*Sw/Hit*) and correct non-switch (*NoSw/Hit*) trials across all recording sites within each brain region. dmPFC and STN LFP recordings were analyzed separately since these data came from distinct groups of patients (dmPFC: n=33; STN: n=24 recording sites). There was a significant increase in theta activity at rule onset when switching, both in the dmPFC (*Sw/Hit vs NoSw/Hit*, [-156 ms; +664 ms], sum(t_(32)_)=232.7, p_c_ <10^−3^, Fig. 2a-b) and in the STN ([4 ms; +1088 ms], sum(t_(23)_)=653.5, p_c_ <10^−3^, Fig. 3a-b), as well as an increase in high-gamma activity (dmPFC: [+98 ms; +625 ms], sum(t_(32)_)=488.4, p_c_ <10^−3^, Fig. 2a; STN: [-523 ms; +189 ms], sum(t_(23)_)=327.8, p_c_ <10^−3^, Fig. 3a). These switch-related changes in theta and high-gamma activity replicated when time-locking iEEG activity at response, which further revealed an increase in dmPFC’s beta (15-30 Hz) activity (*Sw/Hit vs NoSw/Hit*; [-631 ms; 229 ms], sum(t_(23)_)=743.6; p_c_ <10^−3^, Fig. 2a and Fig. S3).

**Figure 2.**
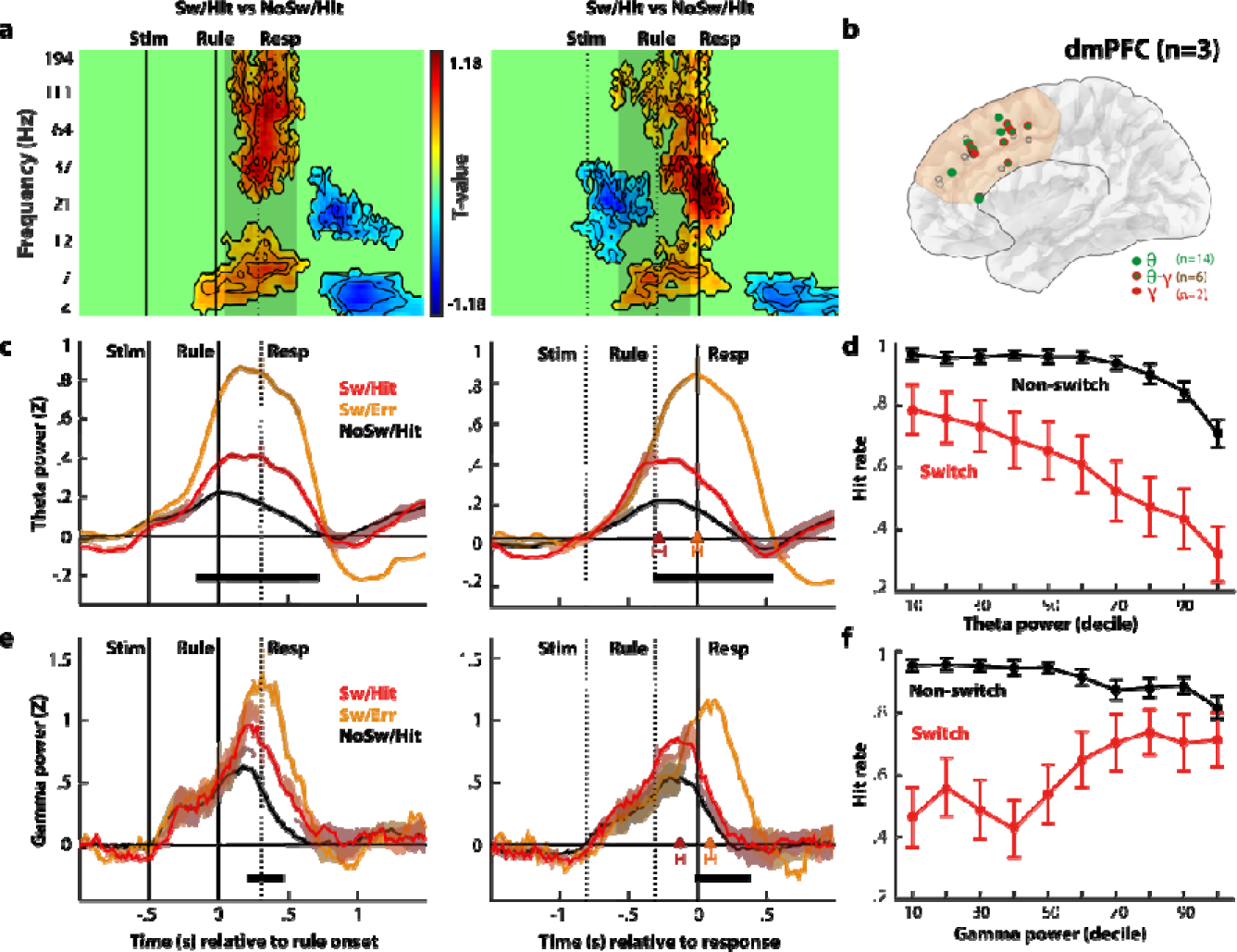
dmPFC neural activity during switching. *(a)* Time-frequency analysis of neural activity time locked on rule (left panels) or on response (right panels) onset across all dmPFC contact-pairs (n=33). Warm (cold) colors indicate significant increases (decreases) of power (p_c_<0.05, FWE cluster-corrected). *(b)* Anatomical location of recording sites in the dmPFC plotted on a 3D reference brain. Each colored dot represents a contact-pair displaying a significant modulation in the theta (green) and/or high-gamma (red) during task-switching. *(c)* Time course of theta power (5-10 Hz) averaged across correct (red) switch, incorrect switch (orange) or correct non-switch (black) trials (n=14 dmPFC contact-pairs). Bold traces indicate average activity and shaded areas correspond to SEM across contacts. The black horizontal bar at the bottom indicates time points for which the statistical contrast between incorrect and correct switch trials was significant (p_c_<0.05). Vertical grey shaded rectangles correspond to 95 % confidence intervals of RT (left panels) or rule onset (right panels). *(d)* Neuro-psychometric curves depicting the relationship between hit rate and averaged theta power (across contact-pairs) binned into deciles separately during switch and non-switch trials. Error-bars correspond to SEM across contact-pairs. *(e)* Time course of high-gamma power (**60-200 Hz**; n=8 dmPFC contact-pairs). *(f)* Neuro-psychometric curves depicting the relationship between hit rate and averaged high-gamma power (across contact-pairs) binned into deciles separately during switch and non-switch trials.

**Figure 3.**
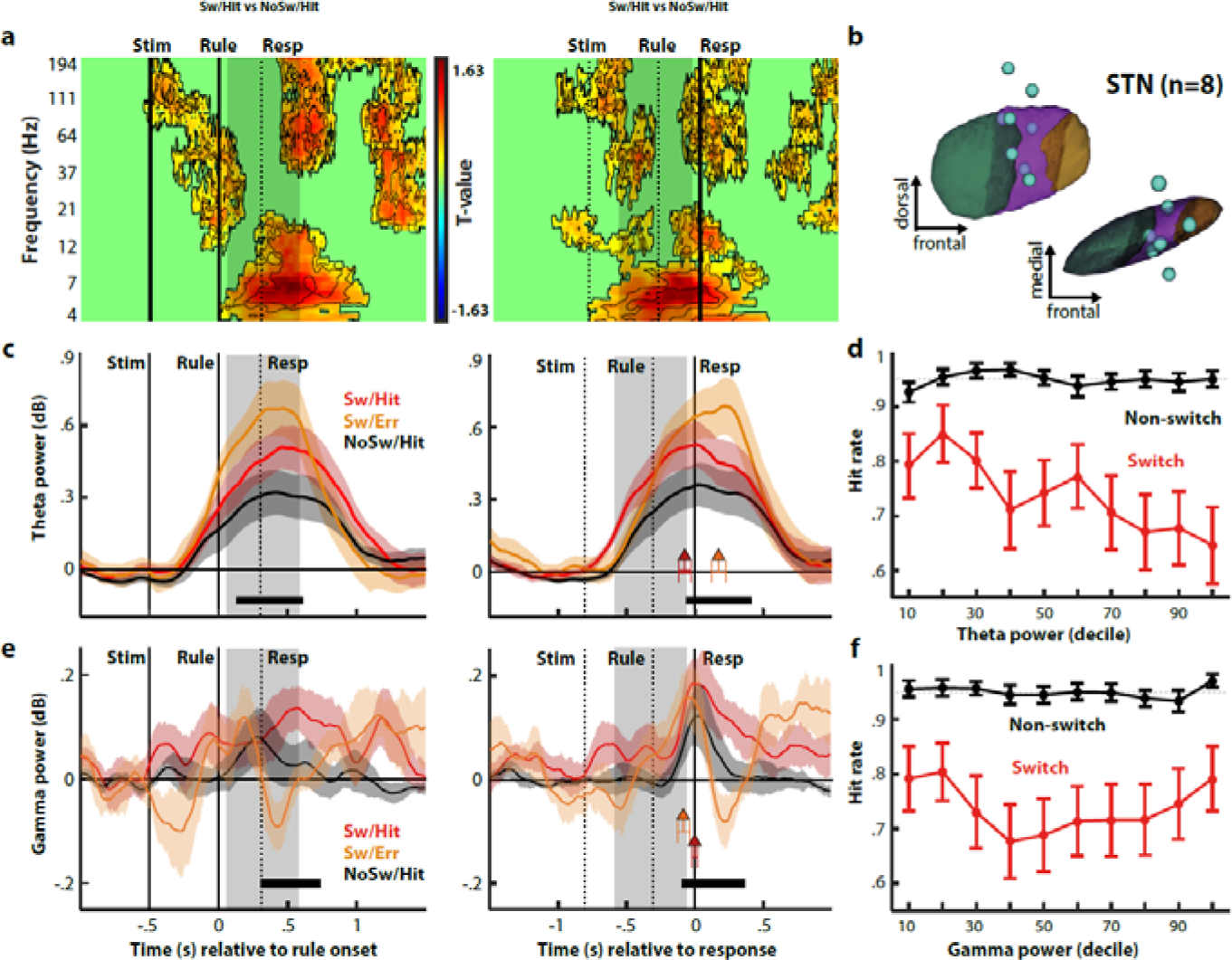
STN neural activity during task-switching. *(**a**)* **Time-frequency analysis** of neural activity time locked on rule (left panels) or on response (right panels) onset (contrasts across all STN contact-pairs; n=24; p_c_<0.05 FWE cluster-corrected). (**b**) **Anatomical location of STN recording locations** with higher task switch related theta increase (blue dots) **displaying switch-related theta activity.** Each contact-pair is plotted on a 3D reconstruction of STN sensorimotor (in green), associative (in purple) and limbic (in brown) territories. ***(c)*Time course of theta power (5-10 Hz) averaged across correct (red) switch, incorrect switch (orange) or correct non-switch (black) trials (**n=8 STN contact-pairs). Bold traces indicate average activity and shaded areas correspond to SEM across contacts. The black horizontal rectangle at the bottom indicates time points for which the statistical contrast between incorrect and correct switch trials was significant (p**_c_**<0.05). Vertical grey shaded rectangles correspond to 95 % confidence intervals of RT (left) or rule onset (right). ***(d)* Neuro-psychometric curves depicting the relationship between hit rate and averaged theta power** (across contact-pairs) binned into deciles separately during switch and non-switch trials. Error-bars correspond to SEM across contact-pairs. ***(e)*Time course of high-gamma power** (60-200 Hz; n=8 dmPFC contact-pairs). ***(f)*. Neuro-psychometric curves depicting the relationship between hit rate and averaged high-gamma power** (across contact-pairs) binned into deciles separately during switch and non-switch trials.

Next, we focused our analyses on contacts exhibiting a differential activity when there was a rule switch or not (i.e., contrasting *switch vs. non-switch trials* independently from accuracy) to investigate the precise time course of theta (5-10 Hz) or gamma (60-200 Hz) band activities in the dmPFC (Fig. 2) and in the STN (Fig. 3). In the dmPFC, we found that the timing of theta and gamma activities differed between switch hit and switch error trials. Theta activity peaked at response when patients failed to switch (0 ± 0.32 ms), but peaked significantly earlier when the switch was successful (*Sw/Hit vs. Sw/Err*, -279 ± 60 ms; t_(13)_= -4.6, p <0.001; paired two-tailed Student’s t test). Consistent with previous neuronal recordings in monkey dmPFC^29^, we observed a similar pattern in the high-gamma band where activity peaked before the response for correct switches (-129±26ms) and after the response when it was missed (+90 ± 40 ms; *Sw/Hit vs. Sw/Err*, -219 ± 42 ms; t_(8)_= -5.23; p<0.001; paired two-tailed Student’s t test, see Fig. 2e). This high-gamma activity peak in the dmPFC reliably followed the peak of theta activity (*Sw/Hit_theta_ vs. Sw/Hit_high_gamma_*, -149 ± 44 ms; t_(21)_= -2.5; p<0.05; unpaired two-tailed Student’s t test).

Furthermore, both theta and gamma activities were also significantly higher for incorrect than for correct switch trials (Theta band: Fig. 2C: *Sw/Hit vs. Sw/Err*, [-299 ms, +522 ms] relative to response, sum(t_(13)_)=85.1, p_c_ <10^−3^; Gamma band: Fig. 2E, *Sw/Hit vs. Sw/Err*, [-6 ms, +366 ms] relative to response, sum(t_(8)_)=57.7, p_c_ <10^−3^).

Taken together, these results demonstrate that successful task switching depends on a structured pattern of increased theta/high-gamma activity in the dmPFC from rule onset to response and precisely controlled both in time and amplitude.

In the STN of OCD patients, similar to what we observed in the dmPFC of epileptic patients, theta activity was significantly higher for incorrect than for correct switch trials (Sw/Hit vs. Sw/Err, see Fig. 3c, [-0.04s, 0.39s] relative to response; sum(t_(7)_)=24.1, p_c_ <10^−3^). Moreover, theta activity in the STN peaked at response (*Sw/Hit* : -0.076±0.042s) when OCD patients successfully switched, but peaked after the response when they failed (*Sw/Err* : +0.172± 0.051 ms; *Sw/Hit vs. Sw/Err, -*219*±*39 ms; t_(7)_= -5.23; p<0.001; paired two-tailed Student’s t test), with a timing (relative to response) reminiscent of the dmPFC’s high gamma activity observed in the group of epileptic patients (*Sw/Hit vs. Sw/Err, -*219*±*39 ms; t_(7)_= -5.23; p<0.001; paired two-tailed Student’s t test). Finally, there was no difference in STN high gamma activity prior to/at the response between correct and incorrect switches, which reflected instead the upcoming motor response (Ipsilateral vs. Contralateral, Fig. S4). Instead, we found that theta increase for successful rule switches was larger in the STN ipsilateral to the newly selected response than in the contralateral STN, but not for unsuccessful switches (see **Fig. S5**, t=2.15; p=0.03), suggesting a direct role for STN theta oscillations in rule-switch execution. Recognizing that dmPFC and STN recordings were obtained from distinct patient groups, thereby precluding dmPFC-STN connectivity analyses, we thought that it remained interesting to compare theta dynamics across these patient groups and structures. Our analysis revealed a consistent temporal lag in rule-switch-related theta increase in the STN compared to the dmPFC (dmPFC-STN theta onsets relative to the rule: dmPFC: -0.419±0.03s; STN: -0.234±0.08s; t_(20)_= -2.57, p=0.018, see Fig. S6). Moreover, the theta increase peaked significantly later in the STN than in the dmPFC (peak latencies in dmPFC: +0.199±0.074s vs. +0.575±0.052s in the STN, t_(20)_= -3.54, p=2.08*10^-3^). Overall, this dmPFC-STN theta/high-gamma dynamic suggests that the dmPFC might drive STN neural activity when external cues trigger a rule-switch and casts the STN as an executive structure downstream the dmPFC along the hyperdirect pathway.

Consistent with this view, we found that neural activity in the dmPFC further encoded the anticipation of upcoming rule switches: dmPFC’s beta activity at stimuli onset increased in proportion with the probability of rule switching (dmPFC: [-846 ms; -534 ms], sum(t_(32)_)=59.99, p_c_ <10^−3^, Fig. S7), 500ms before the onset of the cue indicating that the rule had changed. Indeed, in our task, the probability of switch increased with the number of previous consecutive non-switch trials, and patients implicitly used this information to anticipate switches so that switch costs were large if a switch occurred after 2-3 non-switch trials (switch cost_(2nsw)_ = -43±13%, switch cost_(3nsw)_ = -37±7%) but disappeared if it occurred after 5-6 non-switch trials (switch cost_(5nsw)_ = -17±14%, switch cost_(6nsw)_ = 3.4±13%, see Fig. S7).

Finally, it is noteworthy that our main results were statistically robust and reproduced when using mixed-effect analyses to assess differences in theta activity between correct switch and non-switch trials, both in the dmPFC (t_(5455)_=4.72, p=2.4 × 10^-6^; see also Fig. S8) and in the STN (t_(6244)_=3.69, p=0.00023; see also Fig. S9), as well as between hit and error switch trials (dmPFC: t_(1284)_=4.26, p=2.2 × 10^-5^ ; STN: t_(1332)_=2.33, p=0.019).

### Trial-by-trial fluctuations in theta/high-gamma bands predict successful task switching in dmPFC and in the STN

To explore the functional role of dmPFC-STN theta/high-gamma dynamic, we then investigated whether spontaneous trial-by-trial fluctuations in neural activity related to actual fluctuations in behavioral performance. To do so, we tested brain-behavior correlations between normalized theta or high-gamma activity and performances during switch and non-switch trials separately for each brain region and for each trial type (dmPFC: Fig. 2d-f; STN: Fig. 3d-f).

During switch trials, there was a negative correlation between theta activity trial-by-trial fluctuations and performances, both in the dmPFC and the STN (dmPFC: Fig. 2d; β_hit∼switch_= -0.33 ± 0.06; t_(13)_ = -5.67, p<0.01; STN: Fig. 3d; β_hit∼switch_= -0.13 ± 0.03; t_(7)_ = -4.24, p<0.01). However, during non-switch trials, there was no influence of STN theta activity trial-by-trial fluctuations on performances, while a negative correlation between theta activity and performances was still found in the dmPFC (theta: β_hit∼nonswitch_= -0.14 ± 0.02; t_(13)_ = -6.19, p<0.01), suggesting a functional decoupling of STN and dmPFC theta activities during non-switch trials. Interestingly, we also found significant positive correlations between performances and pre-response high-gamma activity in the dmPFC during switch trials (Fig. 2f; β_hit∼switch_= 0.18 ± 0.05; t_(7)_ = 3.32, p=0.01), whereas this correlation was negative during non-switch trials (Fig. 2f; β_hit∼nonswitch_= -0.09 ± 0.01; t_(7)_ = -5.56, p<0.01). Thus, during non-switch trials, there was a negative correlation, like what was observed in its theta band (and in pre-response beta, see Fig. S10), whereas this correlation became strongly positive during switch trials, suggesting that dmPFC pre-response high gamma activity reflects a neural process selective to task-switching.

### Theta activity is negatively associated with the starting point during rule switching in the dmPFC and in the STN

Finally, to further understand how these trial-by-trial fluctuations in neural activity influenced behavior, we used a hierarchical drift-diffusion model (HDDM^31^) to disentangle behavioral variability arising from switching between rules vs. selecting a new response. At the behavioral level, drift diffusion models (DDMs) are commonly used to model the dynamics of action selection as an accumulation of evidence over time until a certain threshold is reached and a response is triggered^32^ (**see Fig. 4A**). DDMs have four main parameters^26^: the time of non-decision ***t***, the initial level of evidence ***z***, a drift rate ***v*** (the accumulation speed of the information relevant to action selection) and thresholds ***a***, which controls the speed-accuracy tradeoff of the selection process and accounts well for its type I errors^9,28^. In our task, we expected to find a shift of the starting point for switch trials, (1) as correct switches were associated with slower RTs and failed switches with faster RTs (compared to correct non-switch responses; see Fig. 1B), (2) as dmPFC encoded *a priori* the likelihood of upcoming switches and (3) as early behavioral studies suggested that, during task switching, reconfiguring the set of rules seemed to occur before the selection of a new response^26,27^. To test this hypothesis, we first fitted a hierarchical drift diffusion model to RT distributions of correct and incorrect responses from all participants^31^ and tested which model parameter best accounted for observed switch costs (**M1 model space,** also included all possible parameter pairs, **see Methods**).

**Figure 4.**
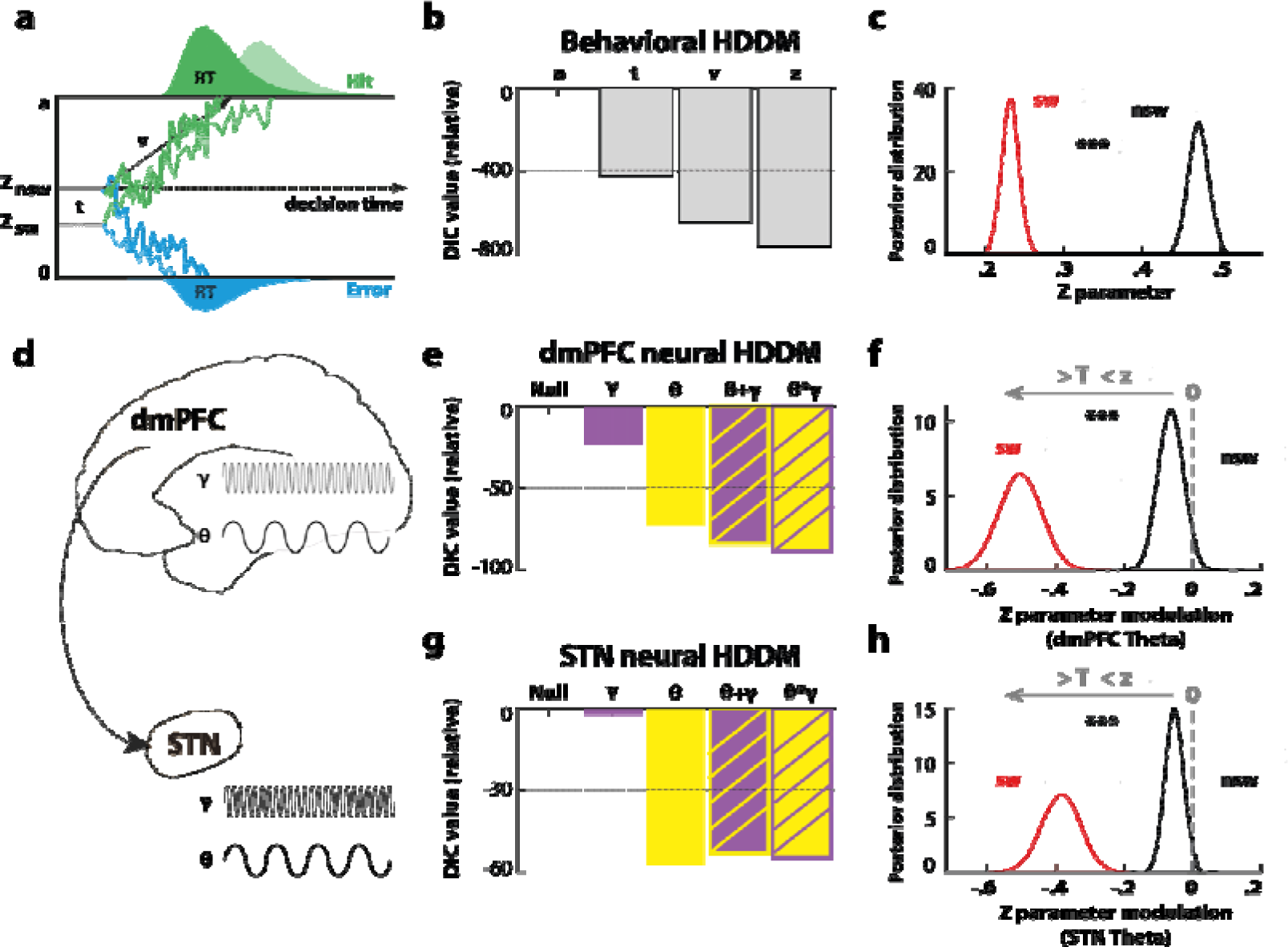
Task switching modeling. *(**a**)*. **Drift diffusion model.** Response execution is preceded by an accumulation of evidence increasing sequentially in favor of one of the two options until a boundary is reached. The decision processes start at stimulus onset and at an initial level depending on the subject’s prior beliefs. The initial level turns out to be different per trial type. *(**b**)*. **Behavioral model comparison.** Relative value of the deviance information criterion (DIC) per model when considering the decision threshold model ***a*** as a reference. *(**c**)*. **Behavioral model posteriors.** Probability density of the initial level of evidence values for switch and non-switch trials. The initial level of evidence is lower during switch trials in accordance with subjects’ belief of an upcoming non-switch trial. The statistical difference between switch and non-switch posteriors was significant (p<0.01). *(**d**)*. **Spontaneous trial-by-trial activities included in the neural HDDM model space. Neural model comparison for dmPFC** *(**e**)* **and STN** *(**g**)*. Relative value of the DIC per model with a model based on a normally distributed noise as a reference. **Neural model posteriors for dmPFC** *(**f**)* **and STN***(**h**)*. Probability density for the effect of theta power on the initial level of evidence z. A negative (resp. positive) regression coefficient means z decreases (resp. increases) when theta power increases. Here, the regression coefficient is negative for non-switch trials and strongly negative for switch trials. The statistical difference between switch and non-switch regression coefficient was significant (p<0.01).

We found that switch costs were best captured by a shift of the starting point (***z*** parameter) toward the lower (erroneous) model boundary (**Fig. 4B**). Moreover, the sampled posterior distribution of ***z*** parameter indicated that HDDM starting point was significantly lower (p<0.05) during switch trials (z_sw_=0.23 ± 0.06) compared to non-switch trials (z_nsw_=0.44 ± 0.06; p<0.001; **Fig. 4C**). Posterior predictive checks confirmed that this HDDM model reproduced all key behavioral patterns observed at rule switch (see Fig. 1B), for each individual in each group (dmPFC Epileptic group; see Fig. S11; STN OCD group; see Fig. S12).

Next, we tested whether dmPFC and STN neural activity further modulated that initial level evidence (***z*)**. We reasoned that using band-specific neural fluctuations to adjust trial-by-trial HDDM’s starting point would significantly improve our model fit only if it reflected the switches between rules. Hence, we built a second neural HDDM model space consisting of HDDMs with starting point modulations by all the switch-related bands identified in our previous analysis of STN and dmPFC neural activity, as well as linear and multiplicative interactions with these frequency bands (**M2 model space,** see Methods, Fig. 4D).

In the dmPFC, the neural HDDM including theta fluctuations as a modulator of the starting point outperformed other single-band models and was not significantly outperformed by more complex neural HDDMs including an interaction term between theta and high-gamma neural activities (see Fig. 4E). Consistent with our other findings in the STN, the neural HDDM model including theta fluctuations as a modulator of the starting point significantly outperformed all the other neural HDDM tested (Fig. 4F). Fig. 4F-H show the posterior distribution of the regression coefficient between HDDM starting point and residual trial-by-trial power in the theta band. The significant shift of the posterior distribution toward negative correlation coefficients between the starting point and theta power (dmPFC: Fig. 4F; STN: Fig. 4H, both p<0.01) is consistent with the observation that, although theta activity in the dmPFC-STN network increases on average during switch trials, too high an increase becomes detrimental to performances.

Some key brain regions, known to exhibit rule-switch activity are not mentioned in this study because an inherent limitation to intracranial EEG recordings is their limited spatial sampling. Note however that the intracranial recordings from the epileptic group allowed us to explore brain activity beyond the hyperdirect dmPFC-STN circuitry. For example, we found that broadband gamma (60-200 Hz) and beta (15-35 Hz) power in the anterior insular cortex was higher during correct switch trials than during non-switch trials (Fig. S13). The main differences with the dmPFC were that (1) there was no modulation of theta band activity in aIns and (2) the increase in both beta and broadband gamma amplitude during incorrect rule switches was only observed after the response, thus lying out of our temporal window of interest (in this study, we focused on the exploration of the cognitive processes related to switch between rules).

## Discussion

In this study, we used human iEEG recordings in either the dmPFC or in the STN to investigate the neural mechanisms associated with rule switching. Our findings reveal new insights into (1) how temporally coordinated neural patterns in the theta/gamma frequency bands in the dmPFC-STN hyperdirect pathway support adaptive rule-switching (2) to the extent that trial-by-trial neural fluctuations of these neural patterns were predictive of fluctuations in observed behavior. (3) We also demonstrate a novel mechanism for rule-switching casting switch costs as a modulation of the starting point in a drift-diffusion process tied, in turn, to a modulation of theta power oscillations in the dmPFC-STN circuit. In the following sections, we will delve into these three lines of results in detail.

### Temporally structured theta-gamma switch-related patterns of neural activity in the dmPFC-STN circuit

By contrasting neural activity between successful trials with and without rule-switching (**Fig. 2-3**), we identified a significant increase of theta and high-gamma activity in the dmPFC-STN circuit between rule onset and response, i.e. when cognitive processes underlying task-switching are theoretically expected to occur.

When contrasting successful and failed switches, we further found that switching accuracy did not only depend on the magnitude of theta and gamma activities: increases of theta (in the STN and dmPFC) and gamma (only in the dmPFC) band activities were also found to peak before the response during successful switches and at the time of the response or later during failed switches (**Fig. 2-3, panels C and E**). Previous monkey electrophysiological studies also reported that neuronal spiking increased in the dmPFC-STN circuit when switching^29,30^, at timings similar to the increase in high-gamma activity observed in our study for successful and failed switches. Taken together, this suggest that a plausible neurophysiological mechanism underlying our findings would be that, in the dmPFC-STN circuit, increased theta band activity may control the timing of the increase in neuronal spiking activity^36^, reflected in our data as an increased activity in the high-gamma just before response. Interestingly, whereas recent iEEG recordings during task switching or stroop paradigms focused on high gamma oscillations^37,38^, these studies did neither investigate the relative timing of theta and gamma activities, nor how these neural proxies relate to behavior. In this study, we found that high-gamma peak activity lagged significantly behind theta’s during successful switch trials, suggesting that dissociable temporal components of switching might be multiplexed by distinct frequency regimes in the dmPFC. An alternative interpretation of the difference of latency observed between theta vs. high gamma peak activity would be that high-gamma increase would partly depend on the preceding theta activity, in line with the extensive literature related to the theta-gamma neural coding schemes underlying several cognitive processes^36^.

The modulation of theta amplitude found in the dmPFC and in the STN during switching is consistent with previous EEG studies in healthy participants^23,25^ and with previous iEEG studies on executive control functions recording LFP across several cortical or subcortical targets^10,13^. Critically, we also found that theta activity magnitude further increased during unsuccessful trials compared to correct switches in both brain structures. Yet, unlike the large body of studies that previously interpreted theta increases during correct and incorrect trials as if these would reflect two independent executive control functions, such as conflict monitoring (in case of success) or error monitoring^18,34,35^ (in case of failure), we propose an alternative view. We identified a single mechanism explaining parsimoniously the pattern of theta oscillations observed in either the dmPFC or the STN: following rule change, theta oscillations amplitude lowers the initial level of evidence available in favor of the new correct response (**Fig. 4**) such that success would only occur for an optimal level of theta oscillations in the circuit.

### Too much or not enough? Trial-by-trial fluctuations of neural activity and behavioral variability

To better understand the respective role of theta and gamma band activities in the prefronto-subthalamic circuit, we examined how the theta and gamma trial-by-trial amplitude fluctuations were associated with behavioral variability across trials, separately for the dmPFC and the STN.

The initial observation was counter-intuitive, since we found that theta activity was negatively associated with hit rate on switch trials both in the dmPFC and the STN. In other words, switching was associated with an increased theta activity observed when contrasting switch vs. non-switch trials, but trial-to-trial analyses revealed that the greater the increase, the lower the switch performance (**Fig. 2D and 3D**). Strikingly, there was no such clear cut relationship between theta trial-by-trial fluctuations and performance for non-switch trials, highlighting its functional dependence to the cognitive context. This finding echoes a previous study showing complex non-linear modulations of decision threshold by STN theta oscillatory activity during simple perceptual decision-making, with a reversal of the relationship as a function of the level of perceived choice difficulty^39^.

At the behavioral level, we found that switch trials (compared to non-switch trials) were best captured by a shift of the starting point (***z*** parameter) toward the lower (erroneous) model boundary in a HDDM^31^. Note that in our task, the erroneous response at switch trials is the response one would have selected if the rule had not been changed. In the context of a reactive rule-switching paradigm inducing reliable increase of RTs when successfully switching between rules, but also premature/fast guess errors when unsuccessful, the fact that the key HDDM parameter associated with task switches was the initial level of evidence is consistent with the idea that subjects were able to prepare the correct response ahead of knowing the rule to be used and, at switch trials, had to inhibit this pre-selected response, switch to an alternative set of rules and then reassess their choice according to it, as previously suggested ^26,27^. Interestingly, we also found that switch probability scaled with pre-rule onset beta activity in the dmPFC (but not in the STN), suggesting that this structure actively monitored and predicted these cognitive events.

At the neural level, we compared several neurally-informed HDDM, in which we tested the benefit of adding either theta or high gamma trial-by-trial activity, or more complex additive or multiplicative models, as factors further modulating HDDM starting point (against no additional neural information). This analysis revealed that across the dmPFC-STN circuit, the most parsimonious mechanism explaining our behavioral results was that trial-by-trial theta fluctuations in this circuit negatively scaled with the starting point parameter of the HDDM. Moreover, this negative correlation was much larger for switch trials (compared to non-switch trials). In other words, when theta power increased at switch trials, it pushed the starting point further toward the lower (erroneous) boundary of the HDDM. This mechanism simply explains how a moderate theta increase allowed patients to buy some time to form their decision when switching to a new set of rules (thus predicting longer correct responses), but also why incorrect switches were more frequent and characterized by very fast reaction times, as more theta activity was observed on average during those trials and its effect on the starting point was amplified, pushing it further toward the lower (erroneous) boundary. The proposed mechanism echoes previous findings from neuroimaging studies in healthy participants suggesting that neural activity in the dmPFC-STN loop may be associated with decision threshold changes captured by drift diffusion models^8,33^ and from studies reporting that applying deep brain stimulation to the subthalamic nucleus of patients with Parkinson’s disease yield complex changes in the regulation of the decision threshold^28,39–42^. To conclude, note that within the theoretical framework of DDMs, only a shift in a starting point can generate such a pattern of RTs combining slower correct and faster erroneous responses for switch trials, as well as an increased error rate (all relative to non-switch trials). Explaining such behavioral patterns is the original motivation behind the introduction of changes (and variability) in DDM’s starting point (Ratcliff et al., 2008). Hence, it is especially meaningful that using trial-by-trial neural variability actually improve the HDDM fits (see Fig. 4e-g).

An obvious limitation of this study is that our data were collected from patients with neuropsychiatric disorders so that disease-related factors could have influenced the neural or cognitive processes of interest. However, it was reassuring to observe that the behavior of patients and healthy subjects was remarkably similar while the neural results were consistent with the results of the animal literature^29,30^. Another limitation is that the dmPFC and STN were recorded in two different groups of subjects, so that direct comparisons between brain regions should be interpreted with caution. That said, differences in pathology or recording modality between the two groups cannot explain the almost identical behavioral and neural patterns we found within each group. The only direct comparison we report pertains to the latencies of theta activity, with earlier onset/peak observed in the dmPFC group compared to the STN group. This is in line with the literature^30^ and with computational models of the cortico-basal ganglia architecture thought to underlie rule-switching^43^. In addition, some key brain regions consistently found to exhibit task-switching signals are absent in our results because an inherent limitation to intracranial EEG recordings is their limited spatial sampling. Yet, we could observe that the anterior insula differentially contributes to behavioral switching compared to the dmPFC (with later timings and a distinct frequency regimes), confirming the interest of intracerebral data to distinguish the functional role of the dmPFC and the anterior insula (Bastin et al., 2017), which are regions often co-activated in functional neuroimaging studies so that their respective functions are still debated in (e.g. Mefford and Critchley, 2010). Examining more precisely the dynamics of task-switching signals in an extended fronto-insular-parietal task-switching network would require additional experiments across a much larger population of implanted epileptic patients performing a task-switching paradigm.

To conclude, we used intracerebral recordings in humans to specify the neural and computational mechanisms through which humans switch between rules: theta activity in the dmPFC-STN circuit adjusts the initial level of evidence such that an optimal level of activity is required to successfully reconfigure the task-set. The existence of non-linear functional associations between neural activities and behavior is important for the development of the next generation of closed-loop neuro-stimulation therapies for neuropsychiatric diseases.

## Methods

### Patients and electrode implantation

The study was approved by local ethics committees (CPP 09-CHUG-12, study 0907 and 2011-A00083-38). *(1) OCD participants.* We included four patients (1 male; mean age 36 ± 3 years old) undergoing bilateral surgical implantation of DBS electrodes with 4 contact leads of 1.5 mm height and 0.5 mm apart (Model 3389; Medtronic, Minneapolis, Minnesota, US) implanted in the STN to treat severe treatment-resistant OCD. These patients were enrolled for STN-DBS according to standard inclusion/exclusion criteria ^44^ and were kept on medication during the experiment (detailed clinical and demographical data are reported in Table S1-a). *(2) Epileptic participants.* Three epileptic patients (no males; mean age 32 ± 2 years old; clinical and demographic details in Table S1-b) suffering from drug-resistant focal epilepsy participated in the study. These patients underwent intracranial cortical recordings by means of stereotactically implanted depth macroelectrodes for therapeutic purposes. All target structures were selected according to clinical considerations independent fromthis study. We recruited patients with at least one electrode in the dorsomedial prefrontal cortex (dmPFC). Nine to twenty semi-rigid macroelectrodes were implanted for each patient. Each macroelectrode had a diameter of 0.8 mm and, depending on the target structure, contained 6-18 contact leads of 2 mm wide and 1.5-4.1 mm apart (Dixi Medical, Besançon, France). All patients volunteered to participate and provided written informed consent prior to participation.

### STN intracranial recordings

We performed intracranial EEG recording from quadripolar DBS electrodes bilaterally implanted in the STNs of four severe and treatment resistant OCD patients using standard clinical procedures (Neurosurgery Department of Grenoble University Hospital) 2–4 days after surgery. LFPs were recorded using temporary lead extensions connected to an EEG acquisition system (Micromed SD MRI, bandwidth 0.15–600 Hz, sampling rate 2048 Hz) before the definitive connection between the electrodes and the stimulation device. Each DBS electrode consisted of 4 contacts with a length of 1.5 mm, separated by 0.5 mm (macro-electrode 3389, Medtronic, Minneapolis, US). We used the tip of the left electrode as reference and computed offline the bipolar derivations between the 3 adjacent pairs of contacts of each electrode to minimize the contribution of sources outside the STN to LFPs. In this study, we only report results based on bipolar derivations because it allows for better signal artifacts removal and achieve high spatial resolution (about 3mm) by canceling out contributions of distant sources, which spread equally to adjacent recording sites.

### STN electrode localization

We reconstructed STN contact locations using a 3D histologic atlas of the basal ganglia warped onto a pre-surgery T1 weighted MRI co-registered to a post-surgery 3D helical TDM [ref_Yelnik_2007, ref_Bardinet_2009]. By analogy with tract tracing data from previous monkey studies ([ref_Yelnik_2007]), this atlas distinguishes 3 functional territories in the STN: posterior/sensorimotor (green in Fig. 2b), intermediate/associative (purple), and anterior/limbic parts (brown). STN contacts shown in fig. 2b are in between the two contacts used for computing the bipolar derivations.

### dmPFC intracranial recordings

Cortical recordings were performed using video-EEG monitoring systems that allowed for simultaneous recording of 128 depth-EEG channels sampled at 512 Hz. Acquisitions were made with the Micromed (Treviso, Italy) system and online band-pass filtering from 0.1 to 200 Hz. Data were acquired using a referential montage with a reference electrode chosen in the white matter. Before analysis, all signals were re-referenced to their nearest neighbor on the same electrode, yielding a bipolar montage.

### dmPFC electrode localization

The electrode contacts were localized and anatomically labeled using the IntrAnat Electrodes software ^45^. Briefly, the pre-operative anatomical MRI (3D T1 contrast) and the post-operative image with the sEEG electrodes (3D T1 MRI or CT scan), obtained for each patient, were co-registered using a rigid-body transformation computed by the Statistical Parametric Mapping 12 (SPM12) ^46^ software. The gray and white matter volumes were segmented from the pre-implantation MRI using Morphologist as included in BrainVisa. The electrode contact positions were computed in the native and MNI referential using the spatial normalization of SPM12 software. Coordinates of recording sites were then computed as the mean of the MNI coordinates of the two contacts composing the bipole. For each patient, cortical parcels were obtained for the MarsAtlas ^47^. Each electrode contact was assigned to the gray or white matter and to an anatomical parcel by taking the most frequent voxel label in a sphere of 3 mm radius around each contact center. Note that data from both hemispheres were collapsed to improve statistical power. Contact-pairs were assigned to the dmPFC region of interest if they belonged to either the caudal dorsomedial PFC or to the rostromedial PFC in the MarsAtlas parcellation scheme.

### Behavioral task

We administered a task-switching paradigm adapted from Isoda and Hikosaka (2010; Fig. 1a). The task included four to five sessions of 200 trials. Each trial started with two colored squares (yellow/pink) randomly displayed on each side (left/right) of a white central square for 500 ms before revealing the ongoing rule (white central square turning either yellow or pink during 200ms to indicate the current target color). Patients indicated their response (left/right with a gamepad Logitech F310S) within a 1000 ms interval by pressing one of two buttons with the index of the corresponding hand. Presentation of visual stimuli and acquisition of behavioral data were performed on a PC (19 inch TFT monitor with a refresh rate of 60 Hz) using presentation software (v14.1; Neurobehavioral systems, Albany, CA). In addition, a time pressure was used to encourage speeded response during non-switch trials; it was initially set to 500ms. This time window was adaptively adjusted by 40 ms steps according to patients’ responses: it was shortened after successful switches, increased after failed switches and could not be inferior to 300 ms. Trials ended with a 1.5s feedback screen displayed immediately after the response or after 1000 ms in the absence of response to indicate the response time and score of the patient using green fonts if the response was correct or a red font if it was incorrect. The score was incremented (decremented) by 1 after each successful (erroneous) non-switch trial and by 5 after each successful (erroneous) switch trial. This procedure ensured that speed and accuracy were balanced: time pressure promoted speeded response while our reward policy promoted high accuracy throughout the task. In between trials, a white square was displayed centrally for fixation (1-1.5s). The rule (yellow/pink) randomly changed after 2 to 6 trials, so that there were 50 switch trials (and 200 non-switch trials) per session. We trained the patient prior to the recording session. First, we provided written instructions and, if necessary, reformulated them orally to ensure they were understood. Then, patients performed a training session of 20 trials (5 switch and 15 non-switch trials) that was eventually followed by a longer training session (maximum 200 trials) to ensure that the patient felt comfortable with the task. The electrophysiological acquisition system was synchronized to the system used to run the behavioral task via TTL pulses signaling the onset of the task events.

### Behavioral analyses

Statistical analyses were performed with Matlab Statistical Toolbox (Matlab R2018a, The MathWorks, Inc., USA). All means are reported along with the standard error of the mean. Trials were excluded from further behavioral and electrophysiological analyses if patients responded before rule onset, if they took more than one second to respond or in the case of response omission.

### Intracranial EEG analyses

iEEG data were first epoched from -3000ms before rule onset to 3000 ms after rule onset. Time-frequency analyses were carried out using the Fieldtrip toolbox for Matlab. Spectral analyses were analyzed in two distinct frequency ranges using a multitaper approach. For the lower frequencies (4-32 Hz), we used adaptive temporal window lengths shifted in steps of twenty ms while fixing the number of cycles (6 cycles and 3 tapers per window. Conversely, for higher frequencies (32-200 Hz), we used a time window length of 187.5 ms shifted in steps of twenty ms with an adaptive number of tappers (4 to 31 tapers : the number of tappers decreased with increasing frequency). This approach uses a constant number of cycles across frequencies up to 32Hz (hence a time window whose duration decreases when frequency increases), and a fixed time window with an increasing number of tapers above 32Hz to obtain more precise power estimates by adaptively increasing smoothing at high frequencies. Spectral power was log-transformed to improve Gaussianity of the data, which allowed us to use standard parametric tests to assess the statistical significance of the observed effects. Theta (5-10 Hz) or high-gamma (60-200 Hz) activity time-courses were extracted by averaging spectral powers (in dB). Theta and gamma activities were normalized (Z-scored) using a [-1500 to -500 ms before rule onset]. We used the full time-frequency analysis to choose upper and lower boundaries for the frequency bands that were identical for dmPFC and for STN recordings, and consistent with previous intracranial studies (e.g., Collomb-clerc et al., 2023; Cecchi et al., 2022; Domenech et al., 2020; Lopez-Persem et al., 2020).

#### Analyses of switch hit compared to non-switch hit trials in the time-frequency domain

(**Fig. 2a and 3a**). For each contact, we computed the unpaired difference in frequency powers at each time step (in the -3000 to 3000 ms relative to rule onset) for every frequency (4-200 Hz) between correct switch vs/ correct non-switch trials. We then computed the corresponding T-values, which we averaged across contacts and participants for each region of interest (dmPFC or STN). Statistical inferences were performed with a corrected for multiple comparison (corrected *p* values, noted *p*_c_<0.05 ; Bonferroni-Holmes Family-Wise-Error corrections at the cluster level; n=150 permutations within contacts, n=50000 permutations between contacts).

#### Selection of task-responsive contact-pairs

For each frequency band of interest (theta or high-gamma), we first selected contact-pairs exhibiting a significant switch-related activity. Switch responsive contact-pairs were identified by (1) extracting the (5-10 Hz) or (60-200 Hz) single-trial power averaged in the time windows during which there was a significant cluster-corrected increase of power time locked to the rule during correct switch trials relative to non-switch trials (dmPFC: [-150 – 560 ms]; STN: [0 – 1100 ms]. We next used these power estimates to contrast for each contact theta or high-gamma power during switch vs. non-switch trials (student unpaired t-test thersolded at p<0.05). All dmPFC contact pairs displaying a significant effect for each frequency band were kept for further analyses because of the spatial sampling heterogeneity inherent to sEEG recordings from epileptic patients. In contrast, all OCD patients were implanted bilaterally within the STN so that we kept one recording site per hemisphere for the STN (choosing the highest T-value among the three contact-pair for each hemisphere), in line with previous human local field potential studies.

#### Analyses of switch error compared to switch hit trials in theta or high-gamma bands

(**Fig. 2c;2e;3c;3e**). For each task-responsive contact, we next computed the unpaired difference in powers at each time step (in the -3000 to 3000 ms relative to rule onset) between incorrect switch vs/ correct switch trials. We then computed the corresponding T-values, which we averaged across contacts and participants for each region of interest (dmPFC or STN). Statistical inferences were performed with a corrected for multiple comparison (corrected *p* values, noted *p*_c_<0.05 ; Bonferroni-Holmes Family-Wise-Error corrections at the cluster level; n=150 permutations within contacts, n=50000 permutations between contacts).

#### Analyses of theta or gamma activity expressed in percentile and hit rate within each trial type

(**Fig. 2d;2f;3d;3f**). For each contact located within our regions of interest, we first sorted theta (or high-gamma) activity into 10 bins. Across the trials within each bin, we then computed for each trial type the accuracy. We then regressed neural activity (expressed in percentiles) and behavioral accuracy separately for each trial type (switch or non-switch trials) for each contact to extract a beta estimate for each contact and each frequency band of interest. We then tested across all contact-pairs within a given ROI (dmPFC or STN) and for each frequency (theta or high-gamma) and for each trial-type (switch or non-switch) whether the regression estimates (slopes) were reliably different from the chance level (two-tailed one-sample t-tests).

### Drift diffusion models

#### HDDM with behavioral data

We fit participant’s choices with the drift diffusion model considering the upper and lower boundaries are accounted for hit and error trials. We used Python 2.7 and the hddm module ^31^ to launch all the model fitting. The first step of our hddm analysis was to modelize behavioural results. For this purpose, we considered each parameter fluctuation as a possible explanation of the difference of reaction time and error rate observed between switch and non-switch trials. Once the DIC compares, it results that a fluctuation of the initial level of evidence is the best explanation of RT and ER difference. We express it as follow:

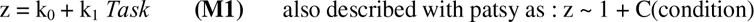

when considering a, t, v constant and independent to task condition (switch or non-switch). Here, k_0_ and k_1_ are coefficients with k_0_ equivalent to the intercept and *Task* refers to the trial type (0 for non-switch, 1 for switch). We generated a chain of 150 000 samples of which the first 100 000 samples were discarded. Thus we obtain a final chain of 50 000 samples and the posterior of all parameters [a,t,v,z] for each subject and for the group. We checked traces of model parameters, their autocorrelation and the model convergence (see supplementary material). The posterior gives information about the probability density for each parameter. So, we can deduce the likelier values and simulate a dataset only based on the modeling (see supplementary material).

#### HDDM with neural dat

In a second step, we analyzed neuro-computational effects during the task. We consider the neural fluctuation through trials and add this information to the previous model. We use the mean value of the power during a specific time window as reflecting the neural state of each trial. We define the initial level of evidence as follow:

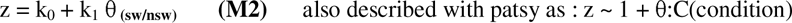

where k_0_ and k_1_ are coefficients, θ the mean power for the frequency range [5-10] Hz. k_0_ acts as an intercept and represents the reference value of the initial level of evidence calculated for all the trials. k_1_ is the coefficient of regression associated with θ which is assigned a certain distribution for switch trials and another for the non-switch trials. Here, we generated a chain of 150 000 samples of which the first 100 000 samples were discarded. Thus, we obtain a chain of 50 000 samples. This time, we calculated the posterior of all parameters for the group only to facilitate model convergence. Indeed, we widely increase the complexity of the model by adding the neural coefficient of regression. To calculate θ value, we used the theta single trial estimation means during the significant increase time window (Fig. 2c and 4c). It gives us a theta estimation per trials which we normalized by subtracting the mean value and dividing by the standard deviation per subject and per trial type^39,42^.

## Supporting information

Supplemental results

## Data and code availability

The data that support the findings of this study and the custom code used to generate the figures and statistics are available from the lead contact (JB) upon request.

## Acknowledgments

This work benefited from University Grenoble Alpes ‘*Investissements d’Avenir*’ program (ANR-17-CE37-0018; ANR-18-CE28-0016; ANR-22-CE17-0057) awarded to JB and from an ‘Agence Nationale de la Recherche’ young investigator grant awarded to PD (ANR-21-CE37-0004-01). The funders had no role in study design, data collection and analysis, decision to publish or preparation of the manuscript. We thank all the patients who participated to the study and Grenoble Teaching Hospital epilepsy, psychiatry and neurosurgery clinical teams for their support. We thank Dr E. Boorman and N. Kolling for their helpful comments on an early version of this manuscript.

## Author Contributions

JB designed the experiment and collected the data. JB, PD, JY, SFV and ML provided preprocessing scripts and anatomical location of dmPFC/STN sites. ML, JB and PD performed the data analysis. MP, PK and SC did the clinical investigation. ML, PD and JB wrote the manuscript. All the authors discussed the results and commented on the manuscript.

## References

1. Monsell, S. Task switching. Trends in Cognitive Sciences 7, 134–140 (2003).

2. Sakai, K. Task Set and Prefrontal Cortex. Annual Review of Neuroscience 31, 219–245 (2008).

3. Isoda, M. & Hikosaka, O. Switching from automatic to controlled action by monkey medial frontal cortex. Nature Neuroscience 10, 240–248 (2007).

4. Isoda, M. & Hikosaka, O. Role for Subthalamic Nucleus Neurons in Switching from Automatic to Controlled Eye Movement. Journal of Neuroscience 28, 7209–7218 (2008).

5. Greenhouse, I., Gould, S., Houser, M. & Aron, A. R. Stimulation of contacts in ventral but not dorsal subthalamic nucleus normalizes response switching in Parkinson’s disease. Neuropsychologia 51, 1302–1309 (2013).

6. Pasquereau, B. & Turner, R. S. A selective role for ventromedial subthalamic nucleus in inhibitory control. eLife 6, (2017).

7. Aron, A. R. & Poldrack, R. A. Cortical and Subcortical Contributions to Stop Signal Response Inhibition: Role of the Subthalamic Nucleus. J. Neurosci. 26, 2424–2433 (2006).

8. Forstmann, B. U. et al. Striatum and pre-SMA facilitate decision-making under time pressure. Proc Natl Acad Sci U S A 105, 17538–17542 (2008).

9. Herz, D. M., Zavala, B. A., Bogacz, R. & Brown, P. Neural Correlates of Decision Thresholds in the Human Subthalamic Nucleus. Current Biology 26, 916–920 (2016).

10. Zavala, B. et al. Cognitive control involves theta power within trials and beta power across trials in the prefrontal-subthalamic network. Brain 141, 3361–3376 (2018).

11. Sheth, S. A. et al. Human dorsal anterior cingulate cortex neurons mediate ongoing behavioural adaptation. Nature 488, 218–221 (2012).

12. Zavala, B. et al. Human Subthalamic Nucleus Theta and Beta Oscillations Entrain Neuronal Firing During Sensorimotor Conflict. Cerebral Cortex bhv244 (2015) doi:10.1093/cercor/bhv244.

13. Kelley, R. et al. A human prefrontal-subthalamic circuit for cognitive control. Brain 141, 205–216 (2018).

14. Smith, E. H. et al. Widespread temporal coding of cognitive control in the human prefrontal cortex. Nat Neurosci 22, 1883–1891 (2019).

15. Kühn, A. A. et al. Event□related beta desynchronization in human subthalamic nucleus correlates with motor performance. Brain 127, 735–746 (2004).

16. Swann, N. et al. Intracranial EEG Reveals a Time- and Frequency-Specific Role for the Right Inferior Frontal Gyrus and Primary Motor Cortex in Stopping Initiated Responses. Journal of Neuroscience 29, 12675–12685 (2009).

17. Swann, N. C. et al. Roles for the pre-supplementary motor area and the right inferior frontal gyrus in stopping action: Electrophysiological responses and functional and structural connectivity. NeuroImage 59, 2860–2870 (2012).

18. Bastin, J. et al. Inhibitory control and error monitoring by human subthalamic neurons. Translational Psychiatry 4, e439 (2014).

19. Benis, D. et al. Subthalamic nucleus activity dissociates proactive and reactive inhibition in patients with Parkinson’s disease. NeuroImage 91, 273–281 (2014).

20. Benis, D. et al. Response inhibition rapidly increases single-neuron responses in the subthalamic nucleus of patients with Parkinson’s disease. Cortex 84, 111–123 (2016).

21. Wessel, J. R., Waller, D. A. & Greenlee, J. D. Non-selective inhibition of inappropriate motor-tendencies during response-conflict by a fronto-subthalamic mechanism. eLife 8, e42959 (2019).

22. Mosher, C. P., Mamelak, A. N., Malekmohammadi, M., Pouratian, N. & Rutishauser, U. *“*Distinct roles of dorsal and ventral subthalamic neurons in action selection and cancellation.” http://biorxiv.org/lookup/doi/10.1101/2020.10.16.342980 (2020) doi:10.1101/2020.10.16.342980.

23. Cooper, P. S. et al. Frontal theta predicts specific cognitive control-induced behavioural changes beyond general reaction time slowing. NeuroImage 189, 130–140 (2019).

24. Proskovec, A. L., Wiesman, A. I. & Wilson, T. W. The strength of alpha and gamma oscillations predicts behavioral switch costs. NeuroImage 188, 274–281 (2019).

25. McKewen, M. et al. Dissociable theta networks underlie the switch and mixing costs during TASK SWITCHING. Hum Brain Mapp 42, 4643–4657 (2021).

26. Schmitz, F. & Voss, A. Decomposing task-switching costs with the diffusion model. Journal of Experimental Psychology: Human Perception and Performance 38, 222–250 (2012).

27. Sinha, N., Brown, J. T. G. & Carpenter, R. H. S. Task Switching as a Two-Stage Decision Process. Journal of Neurophysiology 95, 3146–3153 (2006).

28. Herz, D. M. et al. Distinct mechanisms mediate speed-accuracy adjustments in cortico-subthalamic networks. eLife 6, (2017).

29. Isoda et Hikosaka - 2007 - Switching from automatic to controlled action by m.pdf.

30. Isoda et Hikosaka - 2008 - Role for Subthalamic Nucleus Neurons in Switching .pdf.

31. Wiecki, T. V., Sofer, I. & Frank, M. J. HDDM: Hierarchical Bayesian estimation of the Drift-Diffusion Model in Python. Front. Neuroinform. 7, (2013).

32. Pedersen, M. L., Frank, M. J. & Biele, G. The drift diffusion model as the choice rule in reinforcement learning. Psychon Bull Rev 24, 1234–1251 (2017).

33. Mansfield, E. L., Karayanidis, F., Jamadar, S., Heathcote, A. & Forstmann, B. U. Adjustments of Response Threshold during Task Switching: A Model-Based Functional Magnetic Resonance Imaging Study. Journal of Neuroscience 31, 14688–14692 (2011).

34. Bastin, J. et al. Direct Recordings from Human Anterior Insula Reveal its Leading Role within the Error-Monitoring Network. Cereb. Cortex 27, 1545–1557 (2017).

35. Fu, Z. et al. Single-Neuron Correlates of Error Monitoring and Post-Error Adjustments in Human Medial Frontal Cortex. Neuron 101, 165–177.e5 (2019).

36. Lisman, J. E. & Jensen, O. The Theta-Gamma Neural Code. Neuron 77, 1002–1016 (2013).

37. Tang, H. et al. Cascade of neural processing orchestrates cognitive control in human frontal cortex. eLife 5, e12352 (2016).

38. ter Wal, M., et al. Human stereoEEG recordings reveal network dynamics of decision-making in a rule-switching task. Nat Commun 11, 3075 (2020).

39. Herz, D. M., Zavala, B. A., Bogacz, R. & Brown, P. Neural Correlates of Decision Thresholds in the Human Subthalamic Nucleus. Current Biology 26, 916–920 (2016).

40. Herz, D. M. et al. Mechanisms Underlying Decision-Making as Revealed by Deep-Brain Stimulation in Patients with Parkinson’s Disease. Current Biology 28, 1169–1178.e6 (2018).

41. Green, N. et al. Reduction of Influence of Task Difficulty on Perceptual Decision Making by STN Deep Brain Stimulation. Current Biology 23, 1681–1684 (2013).

42. Cavanagh, J. F. et al. Subthalamic nucleus stimulation reverses mediofrontal influence over decision threshold. Nat Neurosci 14, 1462–1467 (2011).

43. Wiecki, T. V. & Frank, M. J. A computational model of inhibitory control in frontal cortex and basal ganglia. Psychological Review 120, 329–355 (2013).

44. Mallet, L. et al. Subthalamic Nucleus Stimulation in Severe Obsessive–Compulsive Disorder. N Engl J Med 359, 2121–2134 (2008).

45. Deman, P. et al. IntrAnat Electrodes: A Free Database and Visualization Software for Intracranial Electroencephalographic Data Processed for Case and Group Studies. Frontiers in Neuroinformatics 12, (2018).

46. Ashburner, J. Computational anatomy with the SPM software. Magnetic Resonance Imaging 27, 1163–1174 (2009).

47. Auzias, G., Coulon, O. & Brovelli, A. MarsAtlas□: A cortical parcellation atlas for functional mapping: MarsAtlas. Human Brain Mapping 37, 1573–1592 (2016).

