## Supplemental results for "Too much or not enough? Optimal level of human intracranial theta activity for rule-switching in the subthalamo-medio-prefrontal circuit"

**Supplementary materials**

**Table S1: Clinical and demographical data (OCD patients).** M: Male; F: Female; L: Left; R: Right; YBOCS: Yale Brown Obsessive–Compulsive Scale.

| **Patient** | **Sex** | **Age**  **(years)** | **Hand**  **laterality** | **Duration of symptoms**  **(years)** | **Number of STN contact-pairs** | **YBOCS** | **OCD specificity** |
| --- | --- | --- | --- | --- | --- | --- | --- |
| ocd-1 | F | 31 | NA | 24 | 6 | 34 | checking |
| ocd-2 | M | 46 | NA | 26 | 6 | 33 | checking, ording |
| ocd-3 | F | 36 | R | 18 | 6 | 25 | aggressive, contamination, washing |
| ocd-4 | F | 32 | R | 20 | 6 | 30 | checking |

**Table S2: Clinical and demographical data (epileptic patients).** M: Male; F: Female; L: Left; R: Right.

| **Patient** | **Sex** | **Age**  **(years)** | **Hand laterality** | **Duration of symptoms before sEEG**  **(years)** | **Total number of contact-pairs recorded** | **Number of dmPFC bipoles** | **Epileptic Focus** |
| --- | --- | --- | --- | --- | --- | --- | --- |
| epi-1 | F | 30 | L | 20 | 88 | 11 | Right dorsolateral frontal lobe |
| epi-2 | F | 37 | R | 29 | 104 | 9 | Left opercular behind Brocca's area |
| epi-3 | F | 31 | R | 21 | 88 | 13 | Right frontal |

**Table S3: Statistical results performed within each patient on their behavioral performances (RTs and hit rate) and across healthy controls (last raw).** F-value: interaction between task condition (sw vs. nsw) and accuracy (hit vs. error). RTs: reaction times. swh: switch hit; swe: switch error; nswh: non-switch hit; nswe: non switch error. T-values and corresponding p-values: from unpaired two-tailed student tests performed across trials for each patient. HR: hit rate

| **Patient** | **Number of trials**  **(n)** | | | | **ANOVA** | | **SC**  **RT_swh_-RT_nswh_** | | | **SC**  **HR_swh_-HR_nswh_** | | | **RT_swe_ - RT_nswe_** | | |
| --- | --- | --- | --- | --- | --- | --- | --- | --- | --- | --- | --- | --- | --- | --- | --- |
|  | **swh** | **swe** | **nswh** | **nswe** | **F-value (interaction)** | **p-value** | **SC (ms)** | **T-value** | **p-value** | **SC (%)** | **Chi^2^-value** | **p-value** | **RT_swe_ - RT_nswe_ (ms)** | **T-value** | **p-value** |
| ocd-1 | 127 | 65 | 715 | 67 | 29,8 | 6,1E-08 | 86 | 6,4 | 3,2E-10 | -25 | 84 | 4,6E-20 | -59 | -4,42 | 2,1E-05 |
| ocd-2 | 96 | 31 | 567 | 15 | 35,2 | 4,6E-09 | 131 | 7,1 | 3,6E-12 | -22 | 82 | 1,4E-19 | -204 | -3,33 | 0.0018 |
| ocd-3 | 154 | 33 | 748 | 22 | 31,7 | 2,4E-08 | 72 | 6,9 | 8,3E-12 | -15 | 61 | 6,5E-15 | -118 | -4,02 | 1,8E-04 |
| ocd-4 | 121 | 66 | 719 | 59 | 68,2 | 4,9E-16 | 143 | 10,8 | 0,0E+00 | -28 | 103 | 4,0E-24 | -86 | -3,27 | 0.0014 |
| epi-1 | 51 | 47 | 361 | 35 | 7,1 | 8,0E-03 | 70 | 4,8 | 2,1E-06 | -39 | 87 | 1,2E-20 | 2,48 | 0.14 | ns |
| epi-2 | 54 | 41 | 350 | 31 | 6,5 | 1,1E-02 | 84 | 4,8 | 2,3E-06 | -35 | 73 | 1,6E-17 | -1,78 | -0.061 | ns |
| epi-3 | 59 | 23 | 294 | 27 | 23,5 | 1,8E-06 | 106 | 4,6 | 6,8E-06 | -20 | 23 | 1,5E-06 | -133 | -5,39 | 2,1E-06 |
| Healthy subjects (n=10) |  |  |  |  | F(1,36)=15,9 | 7E-04 | 84 ± 12 ms | T(9)=6,8 | 8E-05 | -26 ± 4 % | T(9)=-7,1 | 6E-05 | -60±22 | T(9)=-2,7 | 2E-02 |

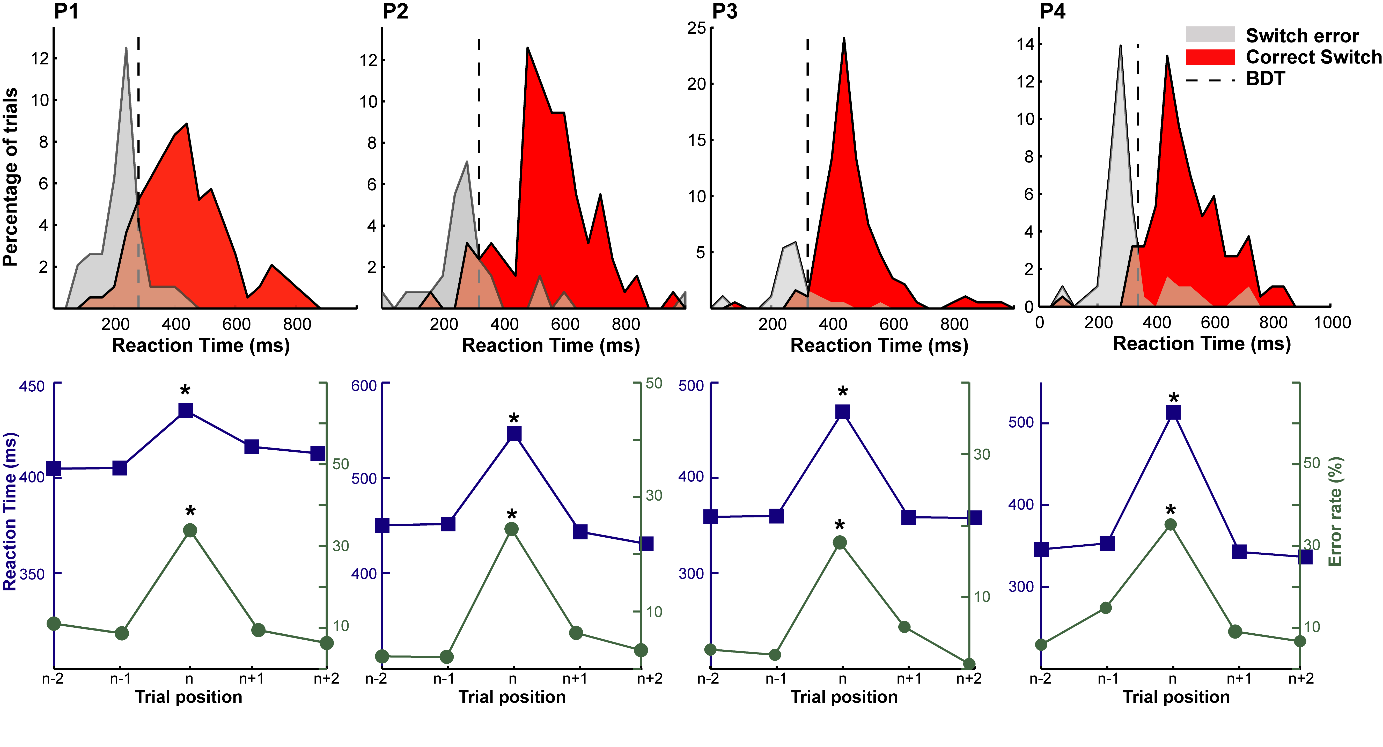

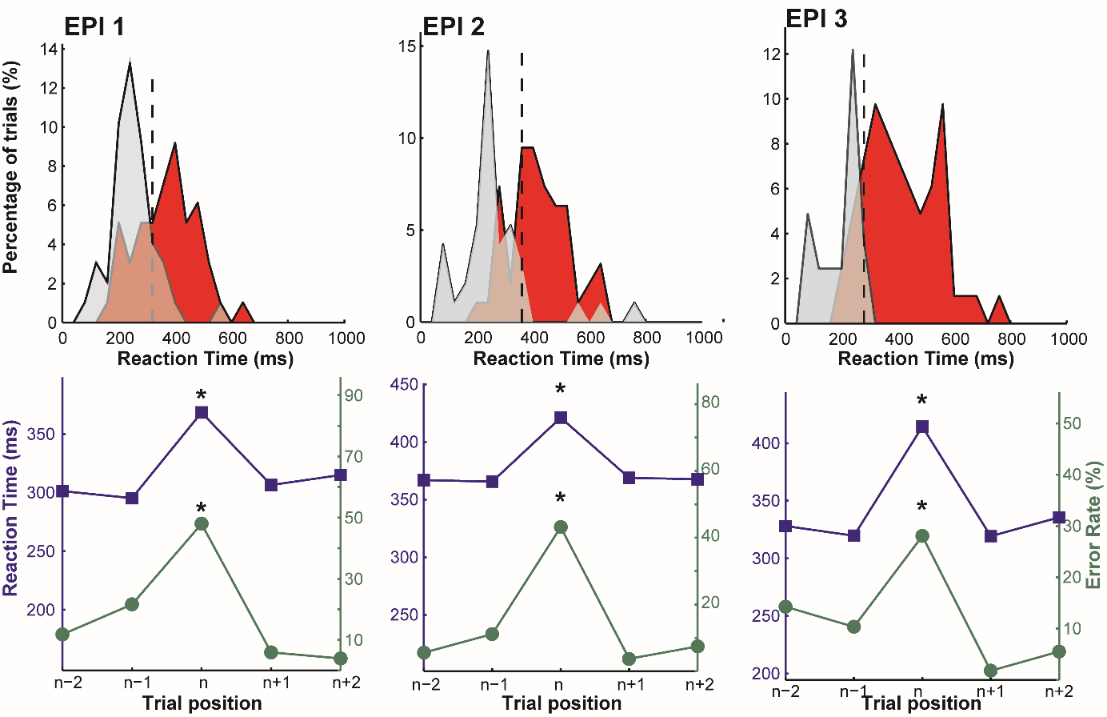

**Figure S1: Behavioral data from the three participants of the dmPFC group**. Upper panels show the distribution of RTs for the switch trials (correct in red vs. incorrect in gray). Black vertical dashed line indicates the behavioral differentiation time between switch errors and switch hit trials (p<0.05, exact Fisher tests for all participants). Lower panels illustrate the evolution of error rates and reaction times averaged according to trial relative position to switch trials (n). Stars indicate switch cost significance (see Supplementary Table X for the individual statistical results on behavior).

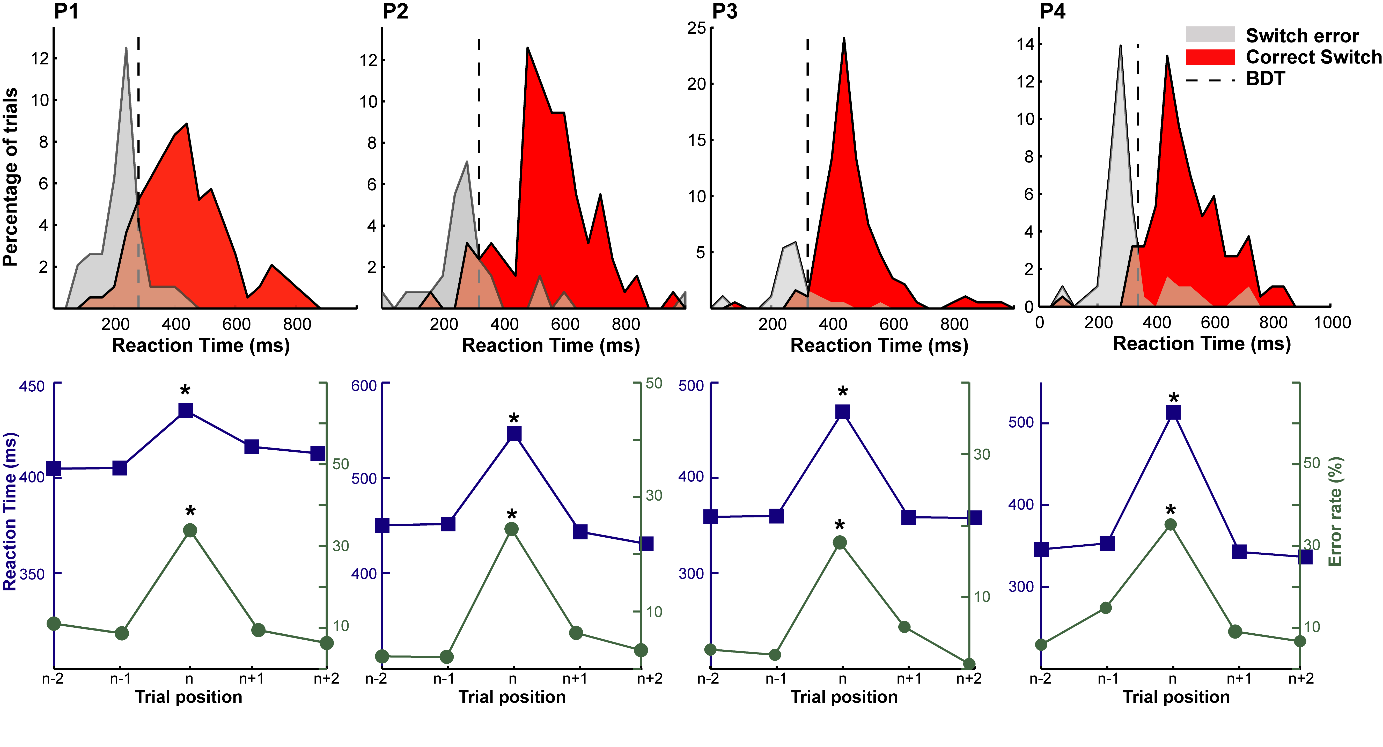

**Figure S2: Behavioral data from individual participants in the STN cohort.** Upper panels show the distribution of RTs for the switch trials (correct in red vs. incorrect in gray). Black vertical dashed line indicates the behavioral differentiation time between switch errors and switch hit trials (p<0.05, exact Fisher tests for all participants). Lower panels illustrate the evolution of error rates and reaction times averaged according to trial relative position to switch trials (n). Stars indicate switch cost significance (see Supplementary Table X for the individual statistical results on behavior).

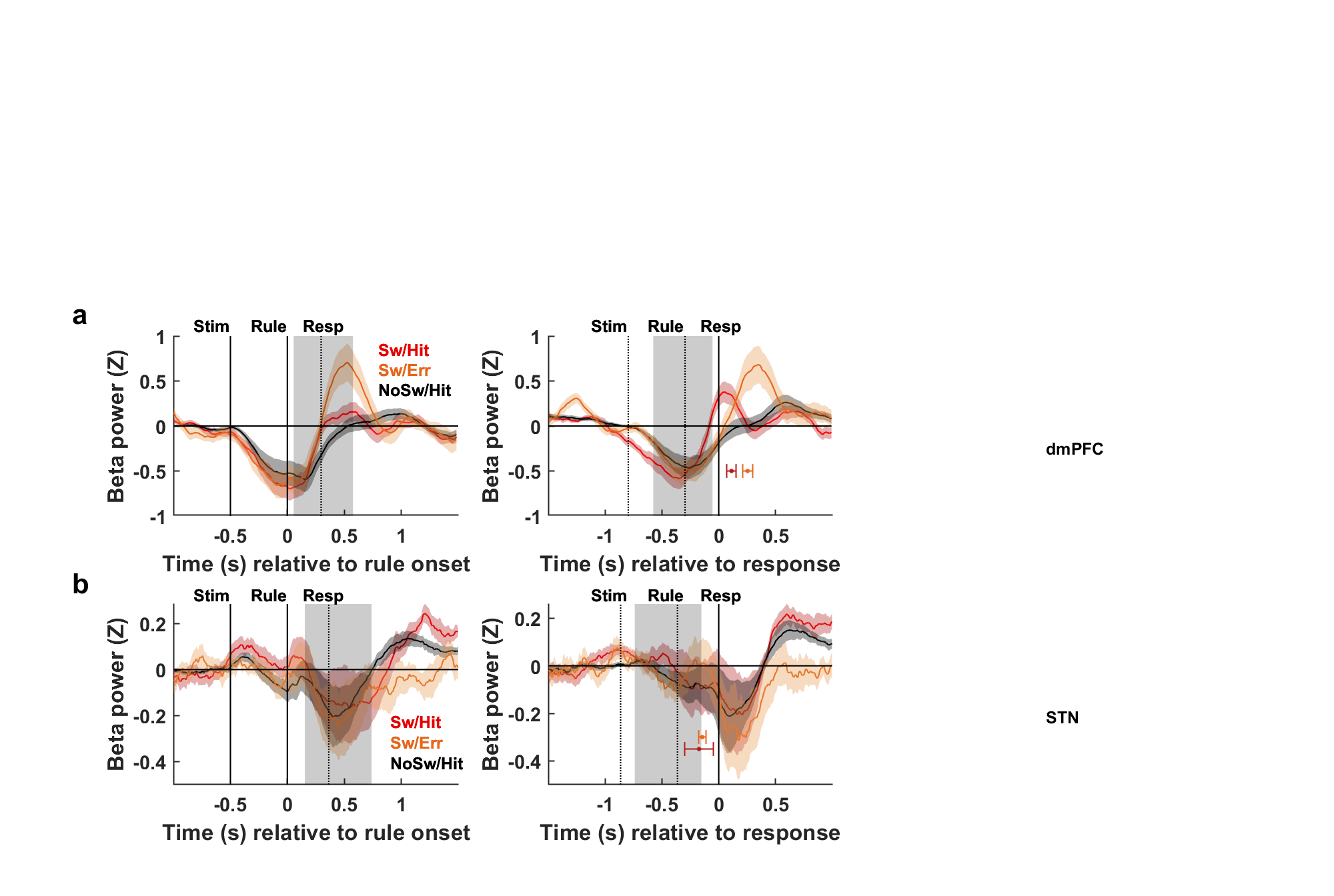

**Figure S3: Neural activity during switching in the beta-band (15-30 Hz) in dmPFC(a) and STN (b). Time course of beta power averaged across correct (red) switch, incorrect switch (orange) or correct non-switch (black) trials** (n=13 dmPFC contact-pairs; n=8 STN contact-pairs displaying a significant beta-band response). Bold traces indicate average activity and shaded areas correspond to SEM across contacts. The black horizontal bar indicates time points for which the statistical contrast between incorrect and correct switch trials was significant (p_c_<0.05). Vertical grey shaded rectangles correspond to 95 % confidence intervals of RT (left panels) or rule onset (right panels).

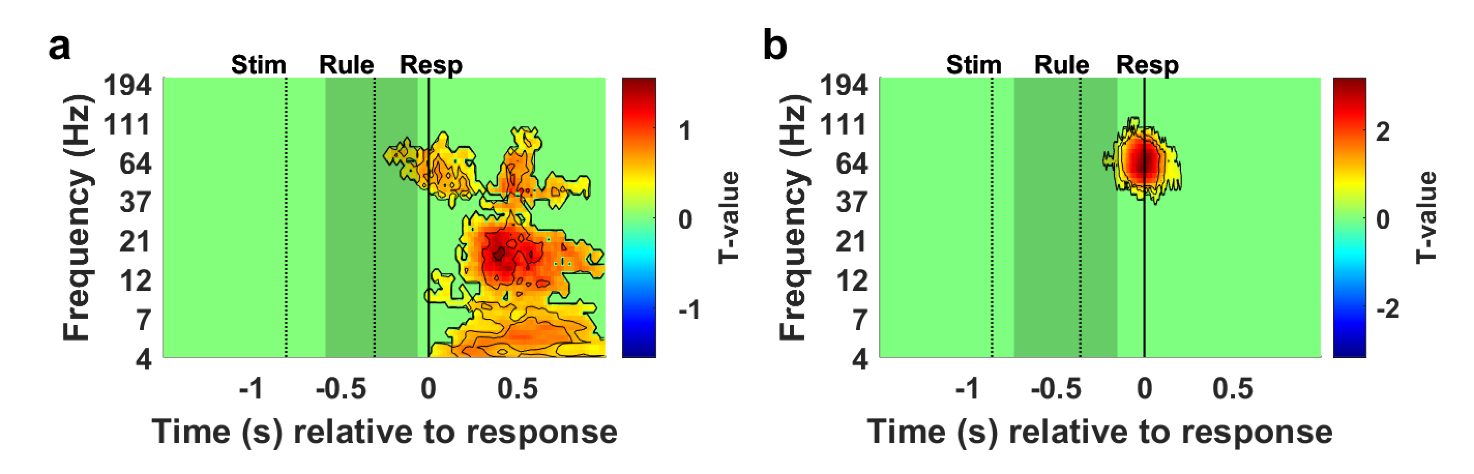

**Figure S4. Time-frequency analysis of neural activity time locked on motor response (each TF map showing the contrast between ipsilateral vs. contralateral responses) across all dmPFC (a) or STN (b) contact-pairs**. Warm (cold) colors indicate significant increases (decreases) of power (p_c_<0.05, FWE cluster-corrected; statistical contrast: ipsilateral – contralateral response). Vertical grey shaded rectangles correspond to 95 % confidence intervals of rule onset.

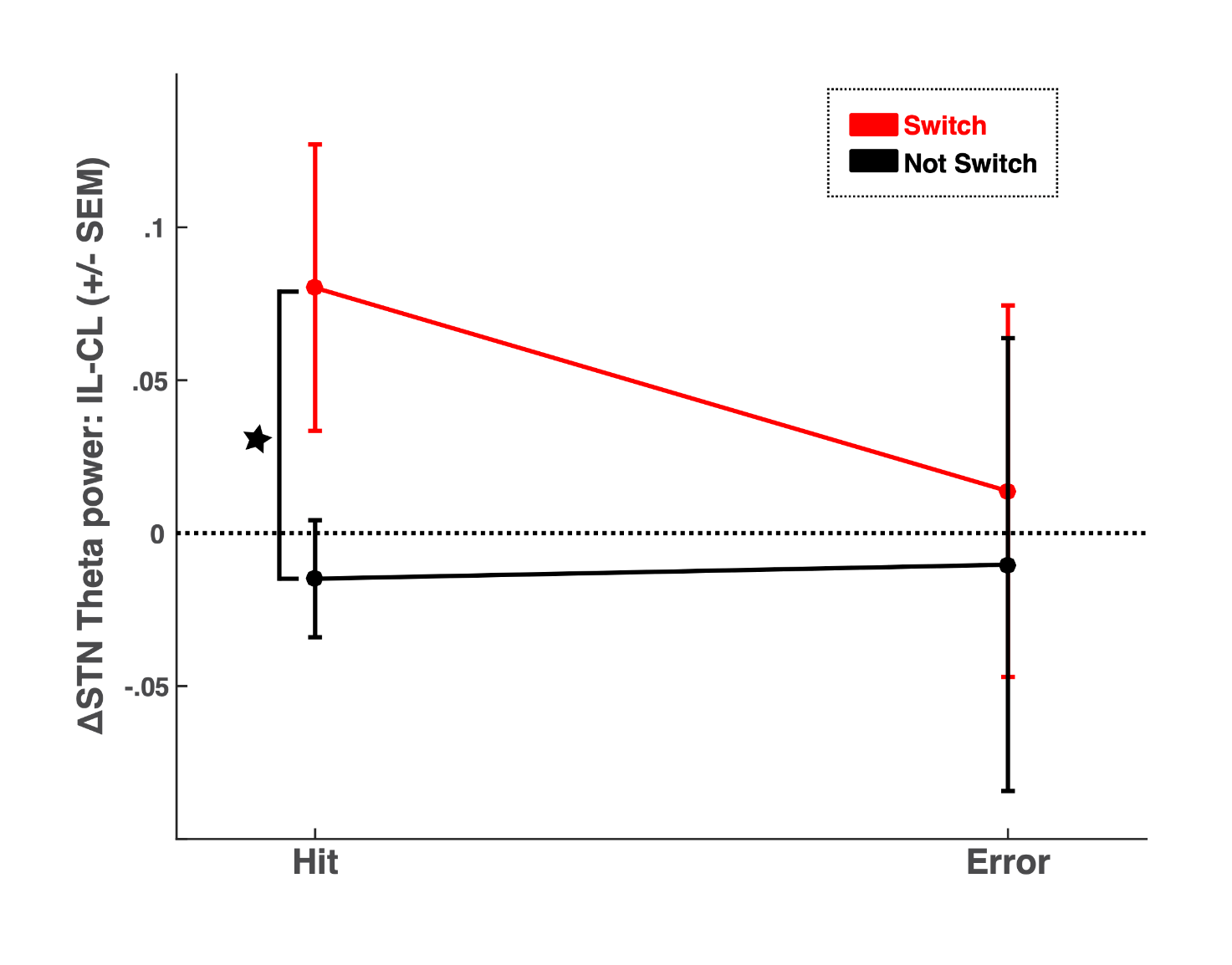

**Figure S5: Theta power increase for successful task switches was larger in the STN ipsilateral to the newly selected response than in the contralateral STN.** Bold dot corresponds to averaged theta power difference between ipsilateral and contralateral theta power across STN contact-pairs (n=8) and error bars represent SEM.

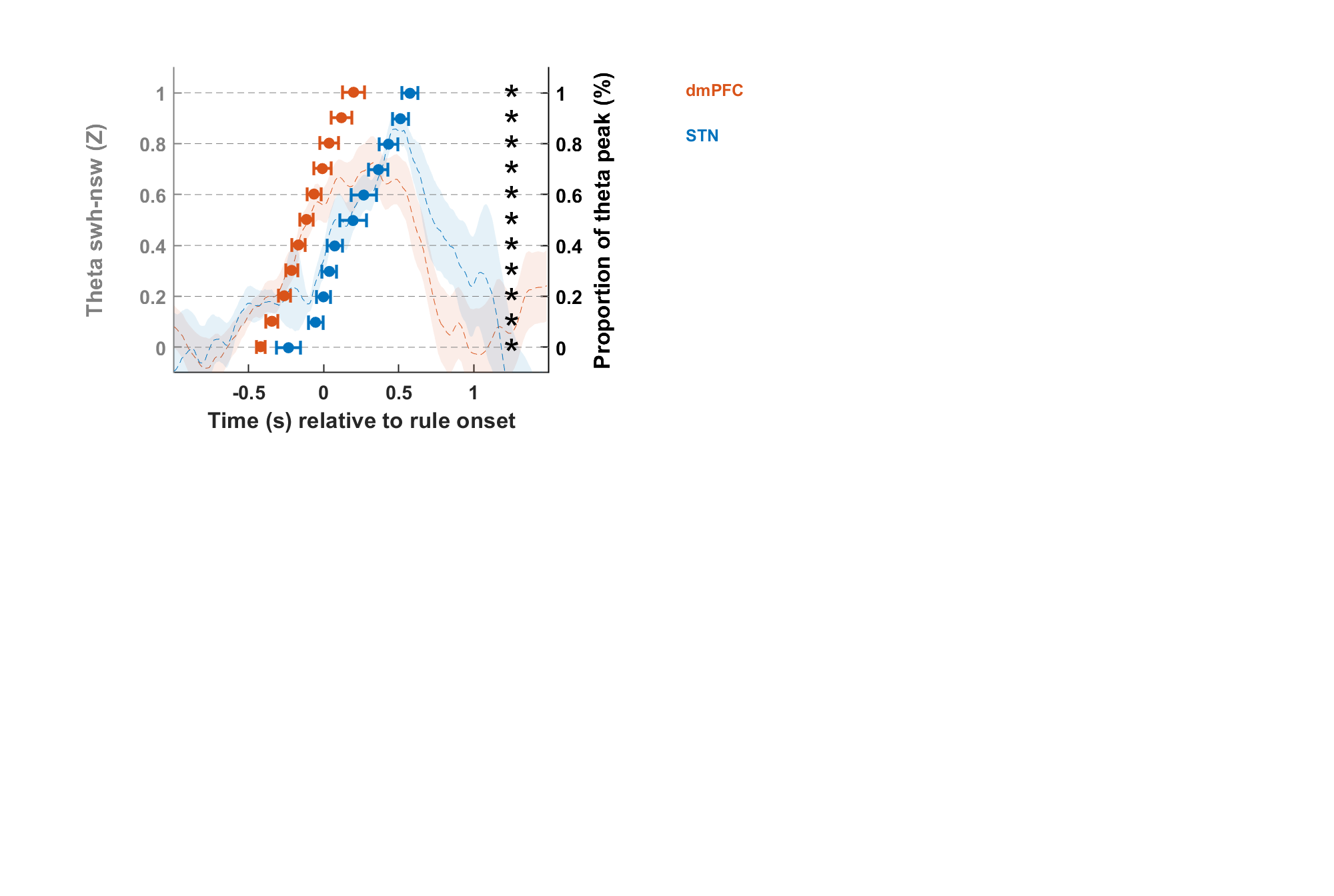

**Figure S6: Theta increase in the dmPFC (red) precedes theta increase in the STN (blue).** Dotted traces indicate the time course of the difference of theta across switch hit and non-switch hit trials averaged across dmPFC or STN contact-pairs (dmPFC: n=14; STN: n=8; Theta (swh-nsw) time-serie of each contact-pair was normalized by Z-scoring the trace using the [-1.5 -0.5 s] time window before the rule as a baseline). Shaded areas correspond to SEM across contacts. To check that dmPFC activity reliably preceded STN activity independently from the threshold chosen to estimate the latency of theta increase, the latency at which theta activity increased was estimated using a wide range of thresholds (proportion of theta increase estimated for each contact-pair maximal theta increase: range [0:0.1:1]). Bold dots and corresponding error bars correspond to average across contacts latency at which theta increased in the dmPFC and in the STN for each threshold (horizontal black dotted lines: 1: theta peak; 0.5: 50% of peak activity). Stars indicate that for all thresholds, dmPFC latency significantly preceded STN latency (unpaired two-sided student t-tests).

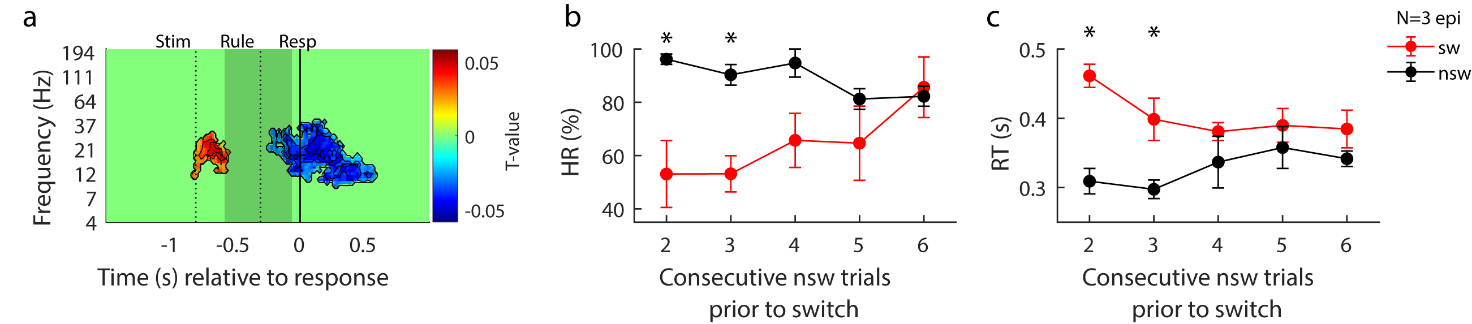
**Figure S7: Neural and behavioral indices associated with the anticipation of upcoming switches in the dmPFC. (a) Time-frequency analysis of neural activity time locked on response onset across all dmPFC contact-pairs**. Warm (cold) colors indicate significant increases (decreases) of power (p_c_<0.05, FWE cluster-corrected; statistical contrast: general linear model between neural activity and the number of non-switch trials preceding the switch (range: 2-6). Vertical grey shaded rectangles correspond to 95 % confidence intervals of rule onset. (b-c) Average across subjects hit rate (HR displayed in panel b) or reaction time (RT displayed in panel c) in the switch and non-switch trials as a function of number of consecutive nsw trial prior rule change. Stars indicate significance (p<0.05 for the difference between SW and NSW trials).

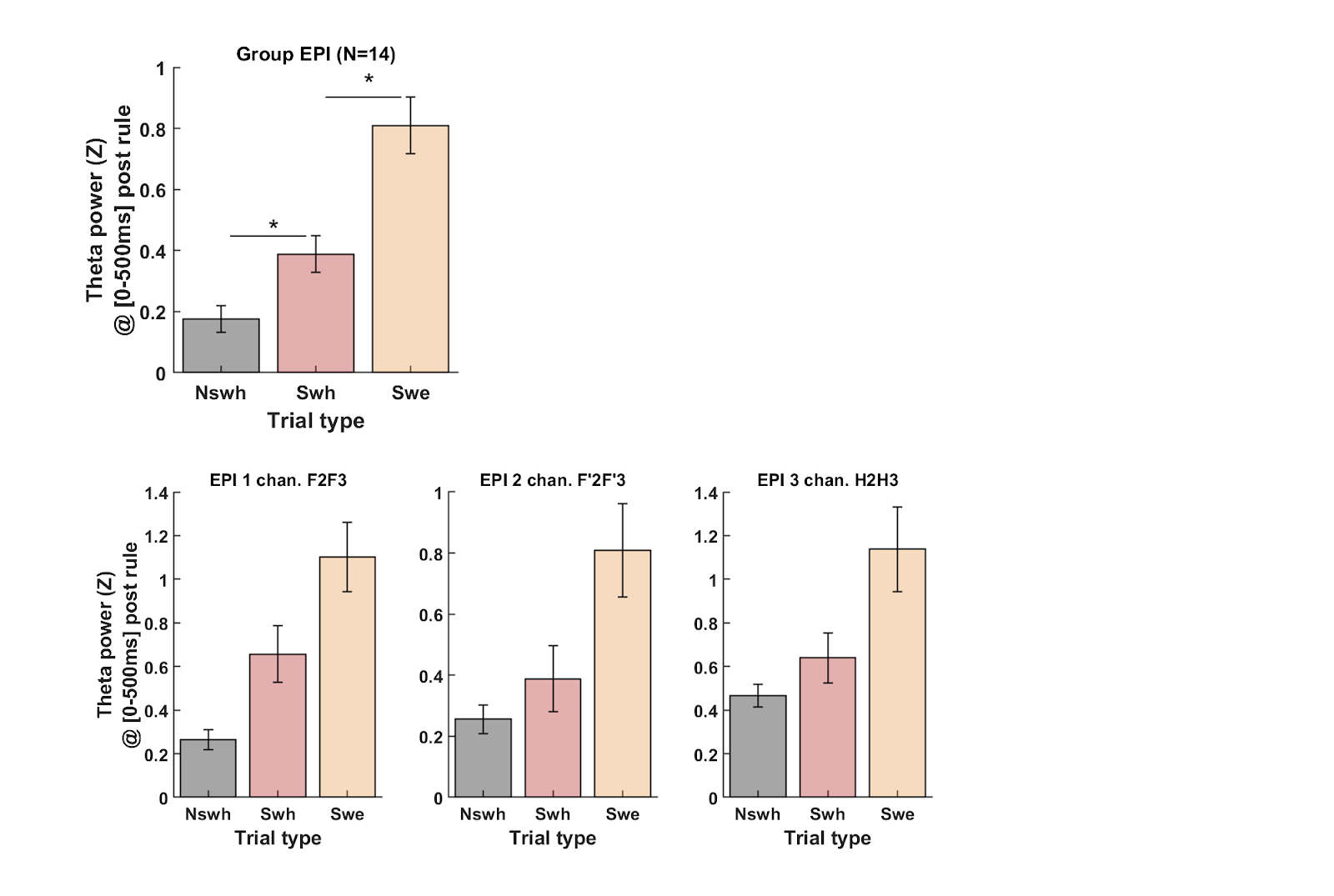

**Figure S8. Linear-mixed effect result (top) and individual pattern of theta amplitude modulation across task conditions in the dmPFC group (bottom).**

Top: For correct trials, we checked using a linear mixed model that theta activity differed between conditions (switch vs. no switch) using the following model applied either on dmPFC data:

TBA ~ 1 + Condition + (1 + Condition | PATIENTS) + (1 + Condition | ELECTRODES:PATIENTS).

This confirmed that theta activity measured in the 500 ms time window post-rule onset was significantly higher for hit switch compared to hit non-switch trials in the dmPFC (t_(5455)_=4.72, p=2.4 × 10^-6^).

For switch trials, we checked that theta activity differed between hit and error trials using the following linear mixed model:

TBA ~ 1 + Accuracy + (1 + Accuracy | PATIENTS) + (1 + Accuracy | ELECTRODES:PATIENTS)

This analysis confirmed that theta activity measured in the 500 ms time window post-rule onset was significantly higher for error compared to hit switch trials either for the dmPFC (t_(1284)_=4.26, p=2.2 × 10^-5^).

Error bars indicate standard error around the mean (SEM) computed across recording sites (n=14).

Bottom: Single recording site theta power across trials shown for each participant. Nsw: non-switch hit trials; Swh: switch hit trials; Swe: switch error trials. Notice that each recording site per subject displayed the group-level pattern. For all three bottom panels, errors bars indicate standard error around the mean (SEM) computed across trials for each recording site shown.

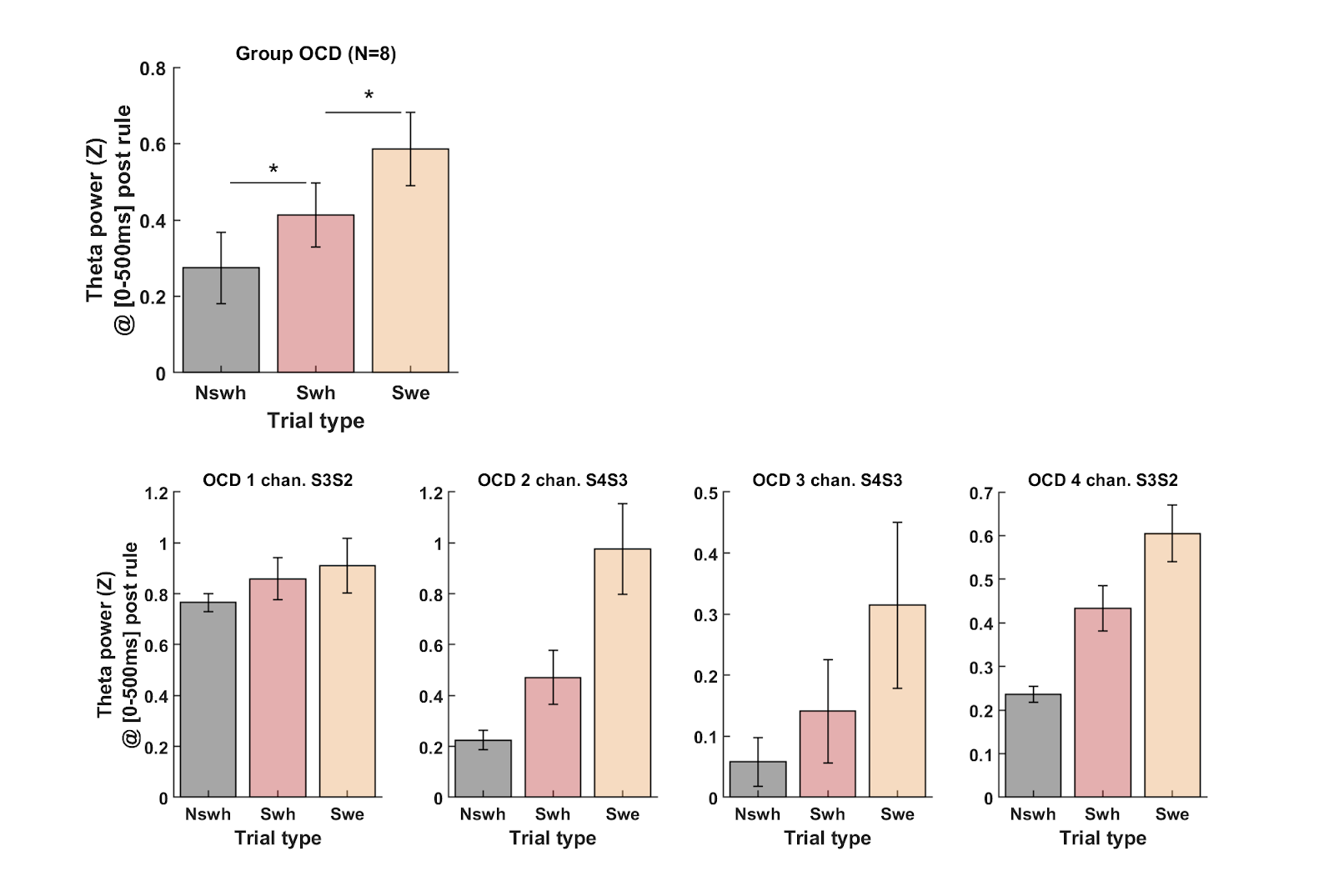

**Figure S9. Linear-mixed effect result (top) and individual pattern of theta amplitude modulation across task conditions in the STN group (bottom).**

Top: For correct trials, we checked using a linear mixed model that theta activity differed between conditions (switch vs. no switch) using the following model applied either on STN data:

TBA ~ 1 + Condition + (1 + Condition | PATIENTS) + (1 + Condition | ELECTRODES:PATIENTS).

This confirmed that theta activity measured in the 500 ms time window post-rule onset was significantly higher for hit switch compared to hit non-switch trials in the STN (t_(6244)_=3.69, p=0.00023).

For switch trials, we checked that theta activity differed between hit and error trials using the following linear mixed model:

TBA ~ 1 + Accuracy + (1 + Accuracy | PATIENTS) + (1 + Accuracy | ELECTRODES:PATIENTS)

This analysis confirmed that theta activity measured in the 500 ms time window post-rule onset was significantly higher for error compared to hit switch trials either for the STN (t_(1332)_=2.33, p=0.019).

Error bars indicate standard error around the mean (SEM) computed across recording sites (n=8).

Bottom: Single recording site theta power across trials shown for each participant. Nsw: non-switch hit trials; Swh: switch hit trials; Swe: switch error trials. Notice that each recording site per subject displayed the group-level pattern. For all three bottom panels, errors bars indicate standard error around the mean (SEM) computed across trials for each recording site shown.

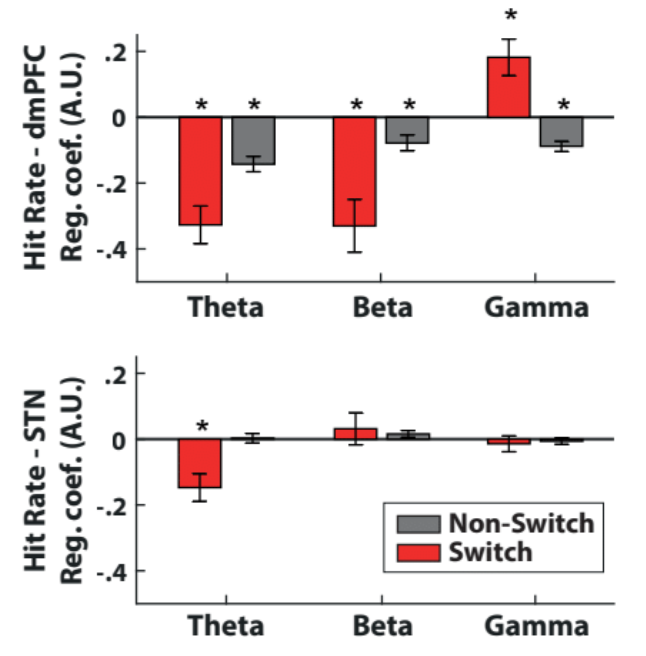

**Figure S10: Association between neural activity and hit rate in dmPFC (a) and STN (b).** Average estimates obtained from the regression of neural activity (averaged within the group-level significant cluster identified before the response in the STN and dmPFC) against hit rate during switch or non-switch trials for each frequency band of interest. Bars represent the average regression estimates across contact-pairs (and error bars represent SEM). Stars indicate significance (two-sided student t-test against chance level).

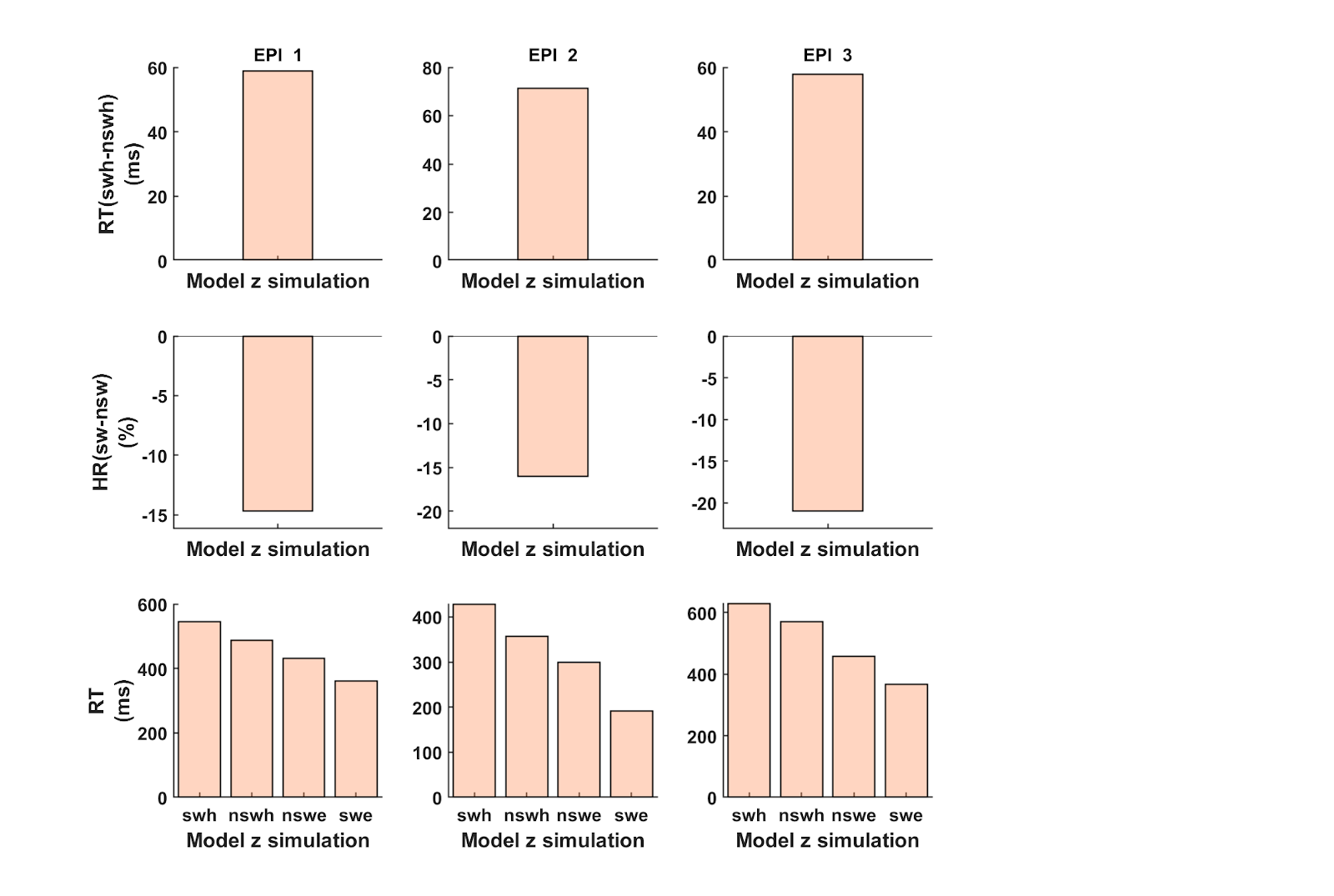

**Figure S11:** DDM model simulation confirmed that a modulation of the starting point predicted increased RT and error rate during switch trials as well as premature responses occurring during errors during switching in all three epileptic participants (dmPFC group).

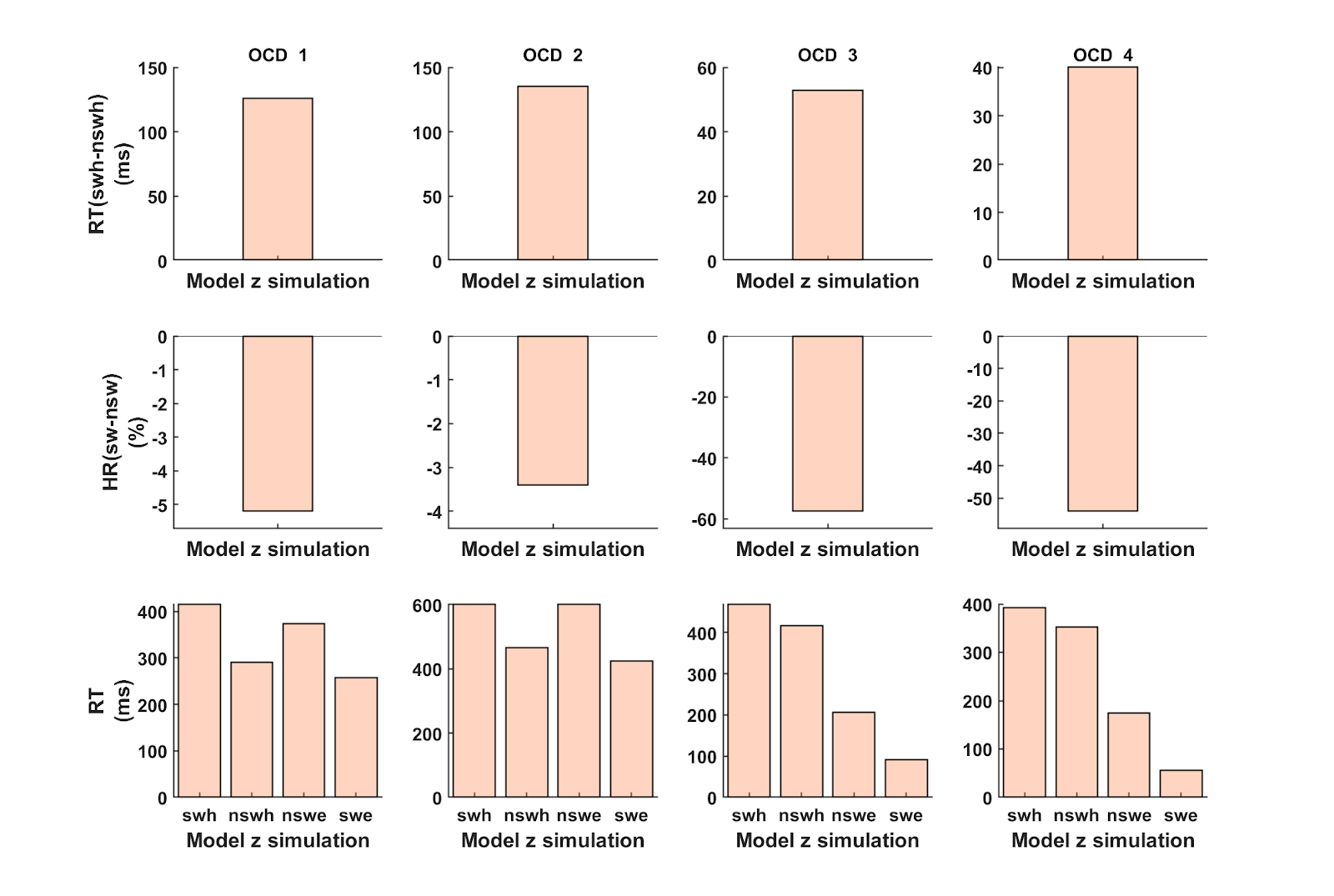

**Figure S12:** DDM model simulation confirmed that a modulation of the starting point predicted increased RT and error rate during switch trials as well as premature responses occurring during errors during switching in all four OCD participants (STN group).

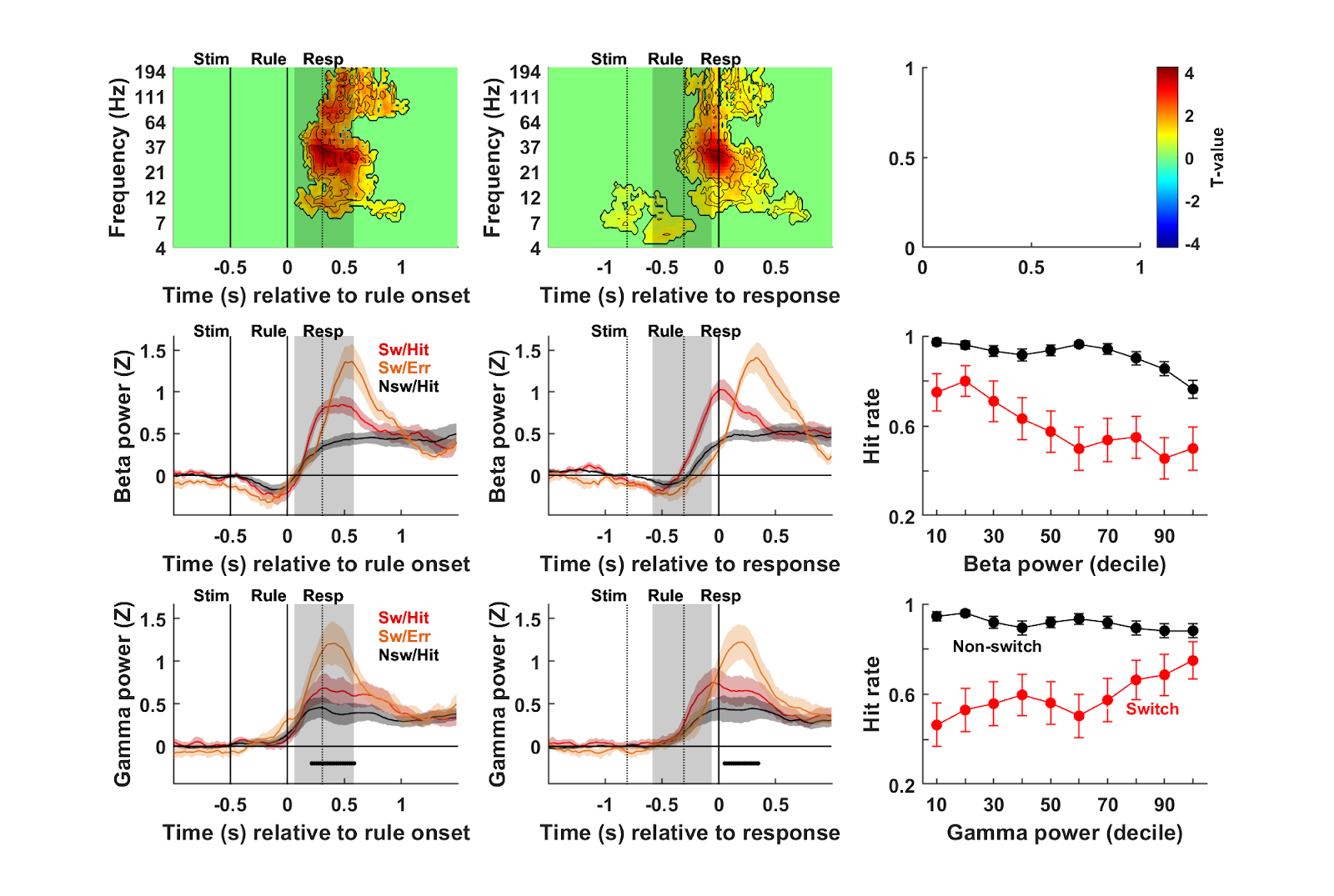

**Figure S13. Neural activity in the anterior insular cortex during switching.** **Time-frequency analysis of neural activity time locked on rule (left panels) or on response (right panels)**. Warm (cold) colors indicate significant increases (decreases) of power. Black contours delimit statistical thresholds from p_c_ < 0.05 to p_c_ < 5.0 × 10^−6,^ FWE cluster-corrected. Significance was assessed using multiple two-sided one-sample student t-tests against zero across all aIns sites (n=20 sites). **Time course of beta power (15-30 Hz) averaged across correct (red) switch, incorrect switch (orange) or correct non-switch (black) trials** (n=9 task-responsive aINS contact-pairs). Bold traces indicate average activity and shaded areas correspond to SEM across contacts. The black horizontal bar at the bottom indicates time points for which the statistical contrast between incorrect and correct switch trials was significant (p_c_<0.05). Vertical gray shaded rectangles correspond to 95 % confidence intervals of RT (left panels) or rule onset (right panels). **Time course of gamma power** (**60-200 Hz**; n=10 task-responsive aIns contact-pairs).
